# Natural genomic variation in rice blast genomes is associated with specific heterochromatin modifications

**DOI:** 10.1101/2023.08.30.555587

**Authors:** David Rowe, Jun Huang, Wei Zhang, Divya Mishra, Katherine Jordan, Barbara Valent, Sanzhen Liu, David E. Cook

## Abstract

Genome organization in eukaryotes exhibits non-random patterns tied to transcription, replication, and chromatin. However, the driving forces across these processes, and their impacts on genome evolution remain unclear. To address this, we analyzed sequence data from 86 *Magnaporthe oryzae* isolates, a globally important plant pathogenic fungus, alongside new reference genomes, to investigate DNA sequence variation and the epigenome. Histone modification profiles were obtained through genome-wide chromatin immunoprecipitation-sequencing of the four new reference strains, which revealed that repressive histone marks (H3K27me3, H3K9me3) were associated with SNP and INDEL frequency. Densely grouped SNPs were found to reside in heterochromatin, often outside transposable elements, highlighting the link between heterochromatin and DNA variation. Even when controlling for selection, silent SNP frequency/kb was higher in H3K27me3-associated genes. Effector genes, key to pathogenicity, also displayed this trend. Comparing the reference strains, euchromatic regions were often syntenic, while heterochromatic regions trended towards non-syntenic. Heterochromatin emerged as a major factor associated with diverse DNA variations in *M. oryzae* populations, even when accounting for selective pressure. This underscores heterochromatin’s pivotal role in shaping genetic diversity in these mainly asexually reproducing fungi.

## INTRODUCTION

Genome variation plays an important role in shaping phenotypic differences within and between species (1–3). There are many forms of genomic variation, two of the most well studied being single nucleotide polymorphisms (SNPs) and small insertions and deletions (INDELs), both of which change short DNA segments. There has long been interest to understand the occurrence of DNA variation within the context of genome organization (4, 5). In fungal plant pathogens, there are contrasting findings regarding the dependence between the distribution and organization of SNPs and INDELs with other genomic features, such as genes and transposable elements (TEs) (6, 7). It therefore remains an open question regarding patterns of DNA variation and the organization of the genome.

The organization of DNA can follow different conceptual frameworks. In fungal plant pathogens, the concept of the two-speed genome has grown in popularity. First described in the oomycete *Phytophthora infestans*, the two-speed genome concept summarizes that densely grouped genes in *P. infestans* have a lower rate of accumulated non-synonymous SNPs, experience less presence/absence variation, and are less transcriptionally induced in the host compared to sparsely grouped genes (8). Often, effector genes, which encode secreted proteins that aid in infection, are organized into similar compartments and regions of the genome (9, 10). However, not all fungal pathogens genomes have a connection between DNA variation and gene density. In the grass powdery mildew pathogen, *Blumeria graminis*, TE insertions have occurred fairly evenly throughout the genome and are found in close proximity to genes regardless of the function of their protein products (11). There are also differences reported in genome organization within families of fungi. In the *Magnaporthaceae* family, an association was observed between unique secreted proteins and DNA variation in *Maganporthe oryzae* (synonym of *Pyricularia oryzae*), but in *M. poae* and *Gaeumannomyces graminis* there was a similar association between DNA variation and genes of any function (12). Taken together, there seems to be a number of recurrent patterns in the organization of fungal genomes, but a generalization has not been described, nor the factors that drive common organization.

The genome is not only organized around DNA sequence characteristics, but also by the epigenome that imparts additional information and regulation. Chemical modifications to histone proteins form an important component of the epigenome (13). Histones form a multi-protein complex, termed nucleosome, that interacts with DNA and provides the local and global structure termed chromatin (14). Chromatin structure, brought about by DNA and protein interactions, plays an important role in many cellular functions including regulating transcription, cellular division, and organismal development (15–18). Histone proteins undergo a variety of post-translational chemical modifications, including methylation and acetylation, which alters their DNA interactions and ultimately their chromatin structure (19–21). Trimethylation on histone H3 lysine 27 (H3K27me3) and lysine 9 (H3K9me3) are considered repressive marks associated with lower transcription, termed heterochromatin (22–25), while methylation of other residues, such as H3K4me3, are associated with active gene transcription and open chromatin termed euchromatin (26). In addition to methylating histones, acetylation of histones can impact DNA usage, such as H3K27ac that is enriched in the promoter regions of transcribed genes (27, 28). Not only do histone modifications regulate transcription, there is increasing interest in how they impact other aspects of genome biology, including evolution (29, 30). For example, in the plant pathogen *Verticillium dahliae*, regions coding for genes involved in plant infection and adaptation are associated with high levels of variation in the population and specific epigenetic profiles (31). In the fungal wheat pathogen *Zymoseptoria tritici*, base substitutions accumulated at higher rates in regions marked with H3K27me3 (32, 33). Additionally, those same regions accumulated base substitutions at lower rates when the gene encoding the histone lysine-methyltransferase were knocked out and the histone mark was absent (33). Interestingly, in *Z. tritici* deletions are more commonly found in H3K27me3 and H3K9me3 marked regions (30), however, in human cell lines, H3K27me3 was negatively associated with the occurrence of new INDELs at double strand breaks (34, 35). A similar pattern can also be observed in *Neurospora crassa*, where H3K9me3 marked regions of the genome have higher mutation rates than euchromatin regions (36). In *M. oryzae*, the accumulation of new SNPs and INDELs in an experimental evolution study occurred more frequently in non-coding regions, and TEs displayed preferential insertion near secreted protein coding genes compared to other types of genes (37). In the model plant *Arabidopsis thaliana*, analysis of mutation accumulation lines showed that genes in euchromatic regions experienced lower mutation accumulation than flanking sequences (38). Interestingly, the observed patterns of mutation largely reflected DNA variation patterns in global *A. thaliana* accessions (38). Together, results across fungal pathogens and in other eukaryotes indicate an association between the epigenome and DNA variation, but the type and direction of the associations, along with the underlying cause, require additional research. A causal understanding of the connection between DNA variation and epigenetic histone marks will provide a deeper understand of genome biology and evolution, and enable new approaches in pathogen control and genome engineering.

To further investigate genome organization in filamentous plant pathogens and its connection with genome variability, we conducted a comparative genomic and epigenomic study in *M. oryzae,* a model plant pathogenic fungus and important global pathogen capable of infecting a wide range of monocots (39–42). We report on highly-contiguous genome assemblies and epigenome maps of four closely related *M. oryzae* rice infecting strains. We show the genomes largely follow organization related to chromatin status, but unexpectedly, all types of DNA variation analyzed, including SNPs, INDELS, gene conservation, and synteny show a strong association with chromatin. The results support the model that fungal plant pathogen genomes exist on a continuum from stable to hyper variable regions, that largely co-occur with epigenome characteristics, and evolutionary analysis indicates that the association may be causative.

## RESULTS

### Complete assemblies of four *M. oryzae* rice-infecting strains have similar coding and repetitive sequence content

To better understand genomic variation between closely related strains of *M. oryzae*, we generated high quality genome assemblies of four rice-infecting strains, named Guy11, O-135, O-137, and O-219. The Guy11 strain is a field isolate collected from French Guiana in 1979 (78), strains O-135 and O-137 are field isolates collected in 1985 from Hangzhou, Zhejiang, China (79), and strain O-219 is a field isolate collected in 1985 from the Ivory Coast (79, 80). Complete genome assemblies were generated using Oxford Nanopore long-read sequencing, ranging from 62.1x to 149.3x coverage (Supplemental Table 1), and further polished with Illumina short-read sequencing. The genomes were assembled into 21-27 contigs each, which were reduced to 8-12 contigs based on quality assessment (see methods for details). Total assembly size for the final assemblies ranged from 42.6 megabases (Mb) to 46.8 Mb with an N50 between 5.7 Mb and 6.5 Mb (Table 1). The contigs were assigned chromosome numbers based on alignment and convention with the 70-15 (50) and B71 (81). Assembly completeness based on Benchmarking Universal Single-Copy Orthologs (BUSCO) (61), which assess the presence of conserved coding sequences, determined that the four strains had similarly high completeness, 97.0%-97.9% (Table 1). Short read genome sequencing data of 82 additional rice-infecting strains of *M. oryzae* were mapped against our Guy11 genome assembly to identify single nucleotide polymorphisms (SNPs). Whole genome SNP phylogeny of the 86 strains formed four distinct clades, consistent with previously published phylogenies of rice-infecting *M. oryzae* strains (41, 82–84) (Figure 1A). Strains Guy11 and O-219 are both present in clade I, which has been described as a diverse group capable of sexual recombination, while O-135 and O-137 represent clades II and III, respectively, both of which are asexually reproducing lineages. Overall, these four strains represent a diverse group of rice-infecting strains.

**Figure 1.**
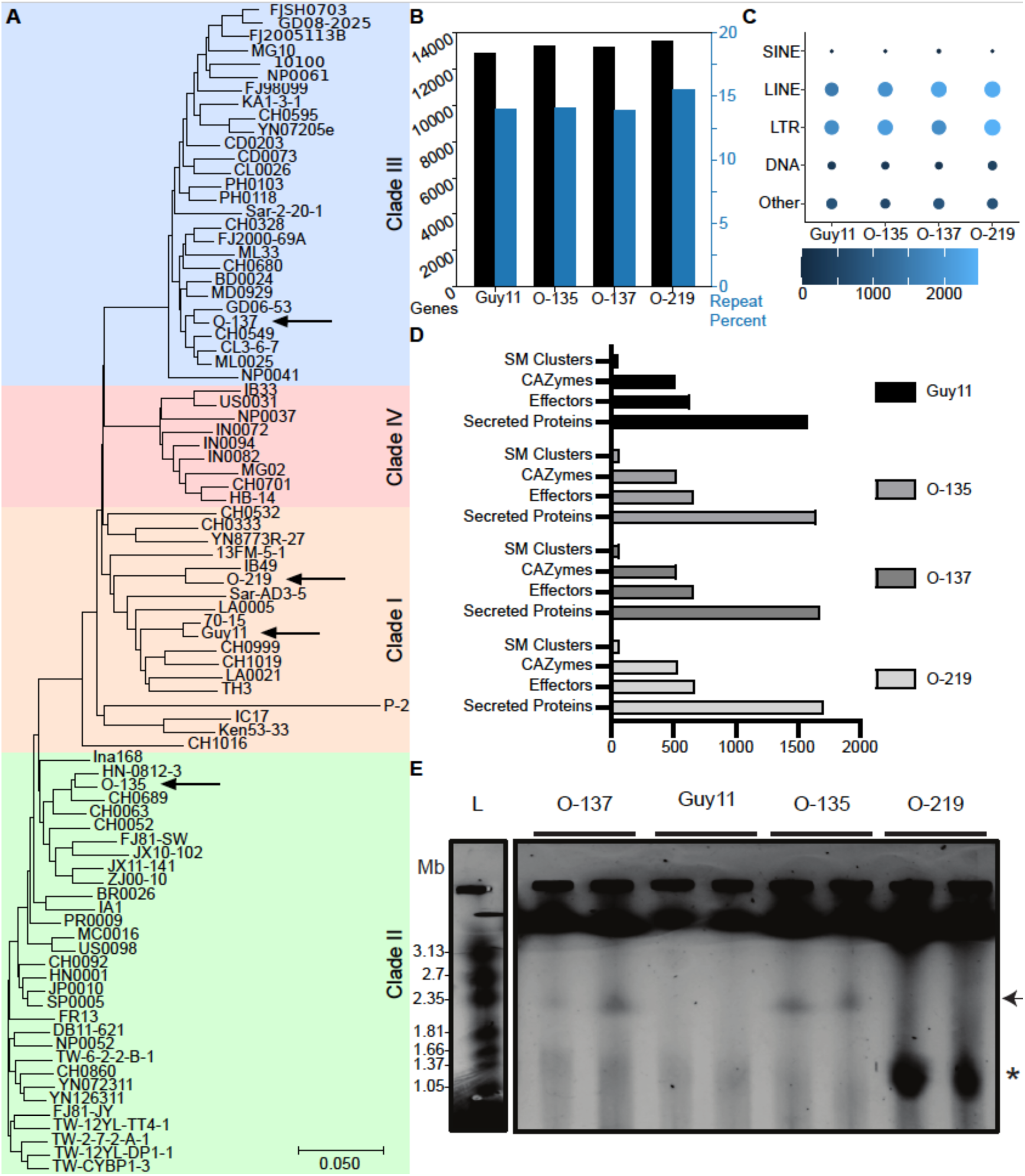
Four rice-infecting *M. oryzae* strain have similar genome compositions based on complete assemblies. (A) Phylogeny of 86 rice-infecting strains of *M. oryzae* using whole genome SNPs compared to Guy11. Clades are labeled to be consistent with previous *M. oryzae* population descriptions (83). Arrows indicate the four newly characterized strains from this work. (B) Total gene count (black) and percent repeat content (blue) of the four strains. (C) Summary counts of TEs separated by class from the four strains. The color bar depicts the number of TEs and the size of each circle is relative to the total count. (D) Counts of secondary metabolite (SM) clusters, predicted CAZyme, predicted effectors, and genes with annotated conventional secretion signals, summarized for each of the four strains. (E) CHEF gel image showing the presence of mini-chromosomes at approximately 2 Mb in strains O-135 and O-137 (arrow), and at approximately 1 Mb in O-219 (star). The assembly indicates multiple minis in O-219 that were not resolved but are indicated by the strong signal.

**Table 1.** Assembly metrics of the four genomes.

| Isolate | Guy11 | O-135 | O-137 | O-219 |
| --- | --- | --- | --- | --- |
| Total long read sequence (Gb) | 3.9 | 5.93 | 2.8 | 6.72 |
| Coverage | 86.6x | 131.9x | 62.1x | 149.3x |
| Contigs | 11 | 11 | 8 | 12 |
| Scaffolds | 2 | 0 | 1 | 1 |
| Assembly size (bp) | 42,642,675 | 44,503,382 | 44,234,998 | 46,802,257 |
| N50 (bp) | 6,590,161 | 5,950,546 | 5,740,312 | 6,382,644 |
| Max contig size (bp) | 8,304,599 | 8,309,243 | 8,325,531 | 8,280,420 |
| Average contig size (bp) | 3,280,206 | 4,045,762 | 4,914,999 | 3,600,174 |
| Genome Completeness | 97.5% | 97.8% | 97.6% | 97.7% |
| tRNAs | 241 | 285 | 303 | 330 |
| Genes | 13,120 | 13,538 | 13,494 | 13,860 |
| Transcripts (mRNA) | 13,050 | 13,341 | 13,344 | 13,462 |
| Secretion Signals | 1,574 | 1,641 | 1,670 | 1,702 |
| Effectors | 634 | 663 | 662 | 670 |
| Protein Completeness | 97.7% | 97.7% | 97.9% | 97% |
| Repeat Content | 14% | 14.1% | 13.9% | 15.5% |
| GC Content | 51.2% | 51.0% | 51.1% | 51.0% |
| Mitochondrial Chromosome | 1 | 1 | 1 | 1 |
| Mating Type | Mat1-2 | Mat1-1 | Mat1-2 | Mat1-2 |

Comparing gene annotation of the assemblies showed that the four strains code for a similar number of genes, ranging from 12,879 to 13,530 (Figure 1B). Plant pathogenic fungi, produce a number of secreted proteins that aid in host infection by subverting or modifying the host immune response (85–88). The presence or absence of functional effectors can strongly influence what plant hosts the fungal pathogen is able to infect (89–91). To identify potentially secreted proteins, SignalP (62) and EffectorP (63) were used to process the annotated protein sequences. Between the four genomes, a range of 1,574 to 1,702 genes were found to have secretion signals (i.e., N-terminal signal peptide) (Figure 1D). Of these, 634 to 670 were predicted to be effectors that may aid infection (Figure 1D). We additionally identified carbohydrate-active enzymes (CAZymes), a group of proteins that catalytically alter or break glycosidic bonds that fungal plant pathogens use to degrade carbohydrates such as components of plant cell walls (92). We identified 519 to 534 CAZymes (Figure 1D). Secondary metabolites (SM) are also commonly produced during host infection, and we identified between 57 and 61 SM gene clusters in the four analyzed genomes (Figure 1D). In addition to protein coding sequences, we annotated transposable elements (TEs) (see methods for details) and found that TEs accounted for 12.5% to 14.4% of the total genome size among the strains (Table 1). Long interspersed nuclear elements (LINEs) and long terminal repeats (LTRs), both of which are retrotransposons, were the two most common classes of TEs, where LINEs accounted for 3.27-4.98% of the genomes and LTR elements represented 6.40-8.31% (Figure 1C). A notable difference between the strains is that three of the four strains were also identified to contain supernumerary ‘mini’ chromosomes. Mini-chromosomes are present in many *M. oryzae* strains but are not necessary for growth (80). O-135 and O-137 genomes each contained one mini-chromosome (Figure 1E). The O-219 genome has previously been reported to contain a large mini-chromosome of approximately ∼2 Mb, and three additional mini-chromosomes on the order of ∼ 1 Mb each (80). Our assembly indicated three mini-chromosomes, ranging in size from 915,411 to 1,401,154 bp, which is consistent with the results from clamped homogenous electric field (CHEF) gel analysis (Figure 1E). Our CHEF analysis was not able to resolve the individual smaller mini-chromosomes, but their approximate size is consistent with the original report and our long-read assemblies. The four assemblies are of high quality and provide new reference genomes for the different clades of *M. oryzae* rice-infecting strains.

### The four strains contain similar numbers of SNPs and INDELs

We next investigated SNPs and INDEL variants in each of the four new assemblies. For SNP analysis, the group of 86 rice-infecting strains were mapped to the four new assemblies and SNPs were called using the Genome Analysis Toolkit (67). The identified SNPs were filtered for biallelic SNPs present in at least 80% of strains for further analysis. The majority of SNPs were found in intergenic regions, averaging approximately 65% (Figure 2A). Of the SNPs occurring in exons of coding sequences, more than half were predicted to be nonsynonymous and cause a change in the resulting amino acid sequence (Figure 2B), similar to previously published data on related strains of *M. oryzae* (93). The INDELs were similarly identified using the 86 rice-infecting strains and Genome Analysis Toolkit pipeline, again using biallelic calls. For this analysis, we defined INDELs as short insertion and deletions that affected DNA sequences between 3 and 30 bp long, while larger identified variants were considered structural variants. Guy11 contained 3265 INDELs, O-135 contained 2495 INDELs, O-137 contained 2661 INDELs, and O-219 contained 2545 INDELs (Table 2). All four genomes had similar percentages of genes containing INDELs, 9.1-9.8%, and similar percentages of TEs containing INDELs, 1.4-2.5% (Figure 2C). The median INDEL size is 4 to 5 bp between the four genomes (Figure 2D). These results suggest that the number and type of short-track DNA variants identified were similar between the four strains.

**Figure 2.**
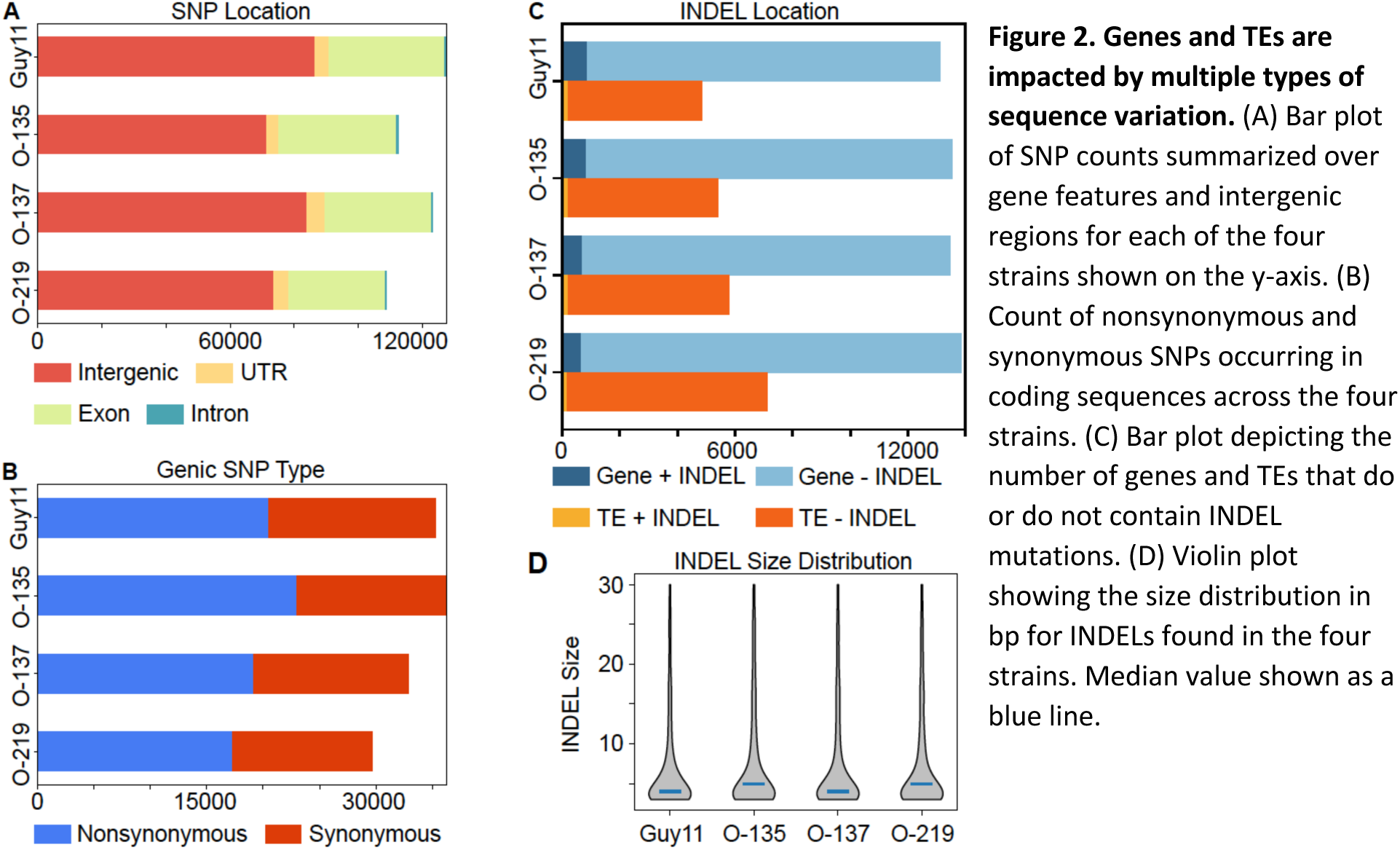
Genes and TEs are impacted by multiple types of sequence variation. (A) Bar plot of SNP counts summarized over gene features and intergenic regions for each of the four strains shown on the y-axis. (B) Count of nonsynonymous and synonymous SNPs occurring in coding sequences across the four strains. (C) Bar plot depicting the number of genes and TEs that do or do not contain INDEL mutations. (D) Violin plot showing the size distribution in bp for INDELs found in the four strains. Median value shown as a blue line.

**Table 2.**
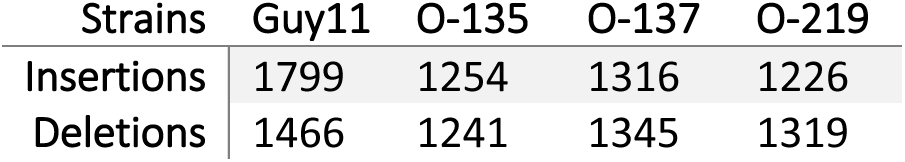
Quantity of INDELs identified in the four strains.

### DNA variants are organized in the genome based on physical location and density

To further understand genome organization, the distribution and density of genomic features were analyzed. Namely, genes, predicted effectors, and TEs, along with DNA variants (SNPs and INDELs) were mapped to their location on the chromosomes (Figure 3). To assess distribution, we divided each of the core chromosomes into ten equally sized bins and summarized the genomic features and variants across all chromosomes of the four strains (Figure 3A, C, E, G, I). Genes were evenly located in the genome across the chromosomes of the four strains (Figure 3A). Each bin represented one tenth of the chromosome, and each bin had approximately 10% of the total genes of any given chromosome (Figure 3A). To assess density, we measured the distance between genes in both the 3’ and 5’ directions. Gene density follows a normal distribution with a 1 kb mean distance between genes, consistent in all four strains (Figure 3B). We also examined the subset of predicted effector genes and found that this specific category of genes was not evenly distributed along chromosomes. Outer bins, representing sub-telomeric sections of the chromosomes, had higher percentages of effectors than inner bins (Figure 3C). The densities of effector coding sequences were not different than that of other genes, occurring approximately 1 kb from each other, and not following a gene sparse pattern (Figure 3D). The TEs were also unevenly located along the chromosomes, similar to genes. Outer bins had on average nearly 20% of the TEs, while central genomic bins contained only 1-5% of total TEs (Figure 3E). The distribution of TE density differs from genes, displaying a bimodal pattern in all four genomes, with a peak of densely occurring TEs within 10 bp of one another, and a second broader peak, with the distance between TEs centered around 1 kb (Figure 3F). This means that some TEs are densely clustered together in the genome. Among eukaryotic genomes, TEs often are inserted near or inside of other TEs which forms nested groups of TEs in repeat-rich regions (94). It is likely that the densely clustered TEs observed in our four genomes are the result of similar nested TE insertions, and also related to difficulty in annotating such complex insertions.

**Figure 3.**
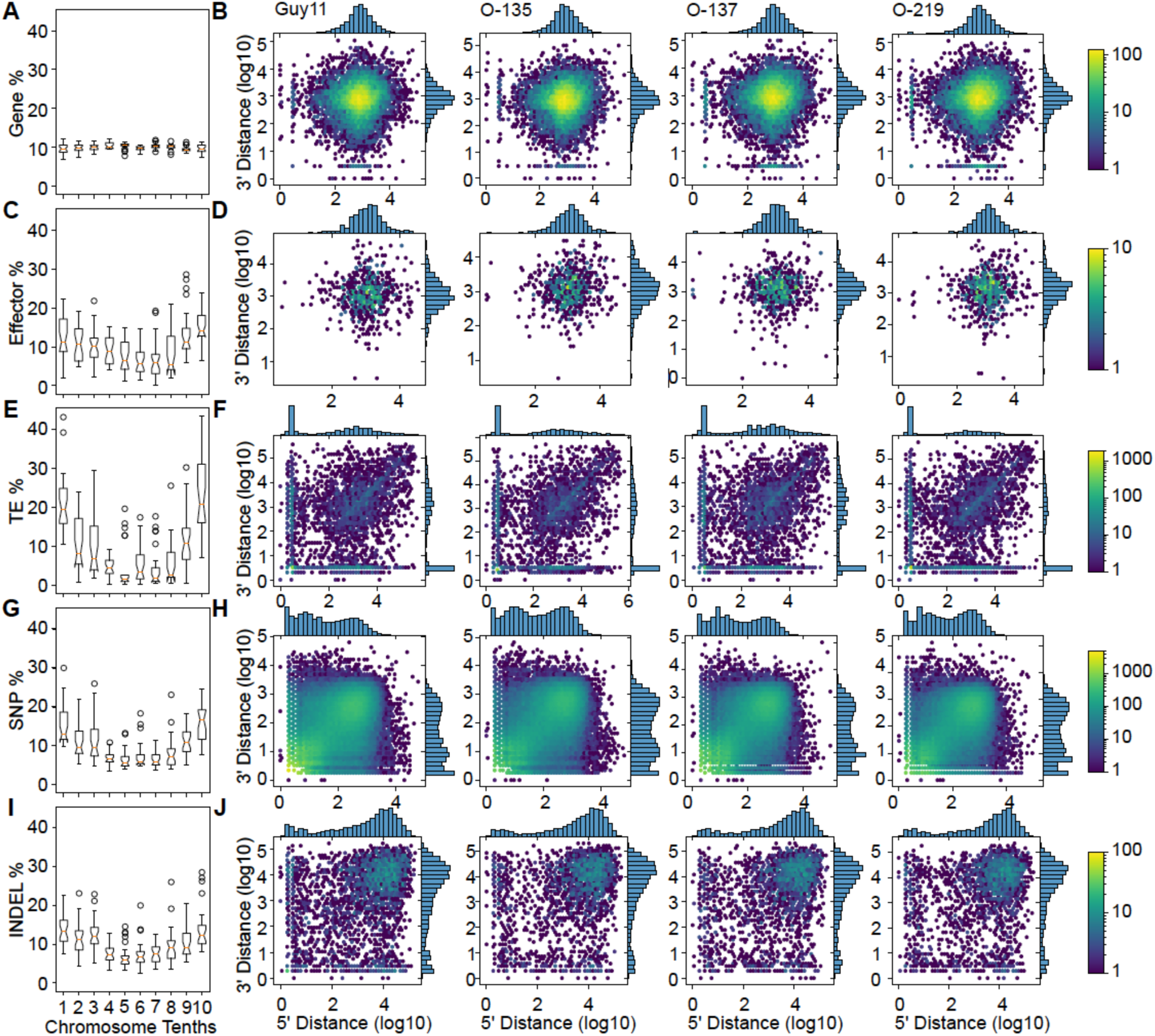
Genomic features display different distributions across the genome from the four analyzed genomes. (A) The percentage of genes grouped into ten bins across the length of each chromosome. Each bin represents 10% of a respective chromosome from each strain. The data are summarized as a box and whisker plot for the same position bin in all core chromosomes across all four strains. (B) Log scaled density plot of gene distribution, representing the genomic distance between each gene in the 5’ (x-axis) and 3’ (y-axis) direction for each of the four strains labeled above each plot. The color bar represents the gene count per hex. (C) The same as A for percentage of effectors across the four strains. (D) The same as B for effector density in each of the four strains. (E) The same as A for percentage of TEs across the four strains. (F) The same as B for percentage of TEs across the four strains. (G) The same as A for percentage of SNPs across the four strains. (H) The same as B for SNP density in each of the four strains. (I) The same as A for percentage of INDELs across the four strains. (J) The same as B for INDEL density in each of the four strains.

Analyzing DNA variants, we found that sub-telomeric regions of chromosomes generally had higher average SNP counts compared to the center of chromosomes (Figure 3G). Interestingly, SNP density forms a multi-modal distribution with a peak around 1 bp, 10 bp, and 1000 bp (Figure 3H). These patterns could also be treated as bimodal, indicating a SNP dense region, defined as SNPs that occur within 100 bp, and a SNP sparse region occurring at a distance of greater than 100 bp. The INDELs also showed an elevated abundance at the chromosome ends, while INDEL density forms a left-skewed distribution with the distance between most INDELs being around 10 kb (Figure 3I, J), indicating that INDELs occur sparsely throughout the genome (Figure 3J). These results show that some functional elements, such as effectors and TEs, and DNA variants, such as SNPs and INDELs are unequally distributed in the genome. The distribution of functional elements and DNA variants as indicated by the density plots indicate common patterns of accumulation between the four analyzed genomes.

### Genome variation is associated with regions containing specific histone modifications

We sought to further understand factors influencing the distribution patterns of functional elements and DNA variants. Previous results indicate an association between genome organization, variation, and the epigenome (31, 33), and we set out to determine whether specific histone modifications are associated with genome organization. Chromatin immunoprecipitation and sequencing (ChIP-seq) data were collected for each of the four *M. oryzae* strains for histone modifications canonically associated with repressed transcription and heterochromatin, H3K9me3 and H3K27me3, and a mark associated with active gene transcription, H3K27ac (13, 95–97). We previously validated the antibodies and experimental protocol for H3K27me3 and H3K27ac in *M. oryzae* (22), and correlation and principal component analysis shows expected and reproducible results from the four strains for the three marks (Supplemental Figures 1-4). Across the four genomes, H3K27ac marked 19.8% to 28.8% of the genomes, H3K27me3 marked 18.8% to 21.1% of the genomes, and H3K9me3 marked 12.4% to 20.7% of the genomes (Figure 4A and Supplemental Figures 5-8). Peak calling of the ChIP data was performed using MACS2 (see methods) to determine regions of the genome enriched for H3K27ac, H3K27me3, and H3K9me3 compared to input controls. Identified peak positions were used to identify overlaps with genes and TEs. Genome wide, H3K27ac marked 13.2% to 22.3% of genes, and H3K27me3 marked 19.7% to 21.7% of genes (Figure 4B). The mark H3K9me3 covered a more variable number of genes between the four genomes, covering 14.0% and 18.9% of genes in Guy11 and O-135, respectively, but only 3.6% and 8.1% of genes in O-137 and O-219 (Figure 4B). As for TEs, H3K27ac covered between 1.2% and 7.3% of TEs, while H3K27me3 marked 36.4% to 54.5% of TEs and H3K9me3 marked 50.6% to 84.4% of TEs (Figure 4C). Given our observation that some SNPs are highly clustered in the genome (Figure 3F), we classified dense SNPs as those within 100 bps of another SNP in both the 5’ and 3’ directions, and all other SNPs were classified as sparse SNPs. Dense SNPs account for nearly half of the total SNPs in the genomes (45.0% to 54.4%), and of those, 20.1% to 33.2% of dense SNPs occur in genes and 6.2% to 45.4% of dense SNPs occur in TEs (Figure 4F). Dense SNPs were intersected with peaks for each of the histone marks, and 3.9% to 8.1% of dense SNPs occur in H3K27ac regions, which stands in contrast to the 38.2% to 58.1% of dense SNPs that occur in H3K27me3 regions and 53.1% to 74.6% of dense SNPs that occur in H3K9me3 regions (Figure 4D). The association between dense SNPs and heterochromatin marks is not simply driven by TEs, as over half of the dense SNPs in each genome occur outside of TEs (Figure 4F). Sparse SNPs on the other hand have moderate association with all measured histone modifications, highest for H3K27ac marked regions (19.5% to 27.6%) (Figure 4E).

**Figure 4.**
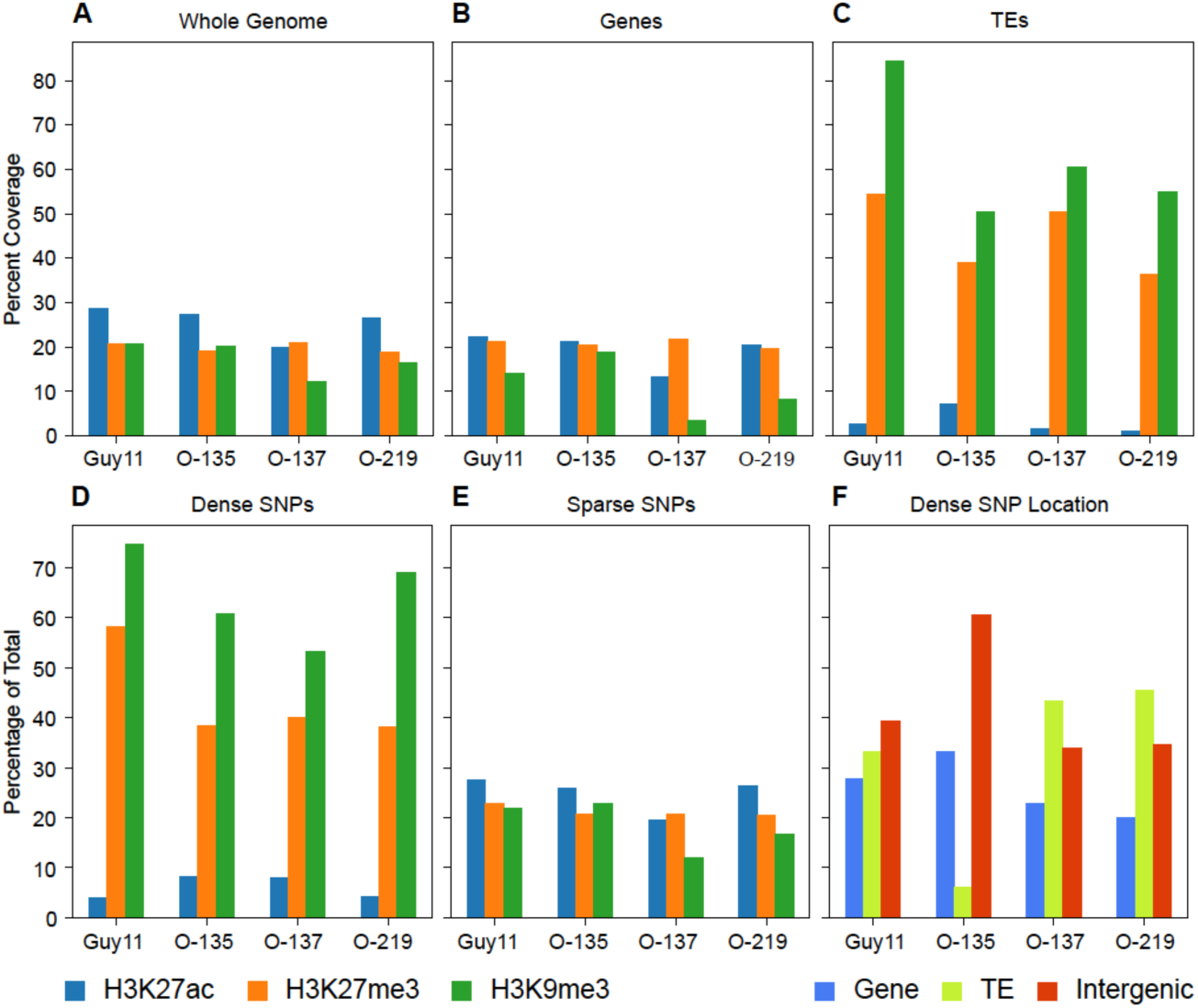
Densely grouped SNPs more frequently occur in repressive histone marks. (A) The percentage of the genomes covered by H3K27ac (blue), H3K27me3 (orange), and H3K9me3 (green) averaged across replicates for each genome shown on the x-axis. (B) The same as A showing percent coverage over genes. (C) The same as A showing percent coverage over TEs. (D) The percentage of dense SNPs, defined as SNPs within <100 bp distance to the nearest SNP in 5’ and 3’ direction, which occur in domains of H3K27ac, H3K27me3, and H3K9me3. (E) The percentage of sparse SNPs, defined as SNPs with >100 bp distance to the nearest SNP in either 5’ or 3’ direction, which occur in domains of H3K27ac, H3K27me3, and H3K9me3. (F) The percentage of total dense SNPs that occur in genes, TEs, and intergenic regions for each strain noted on the x-axis.

To gain a broad understanding of the relationships between the analyzed variables, all the data, including counts of genes, TEs, SNPs, INDELs, and the normalized ChIP signal for each histone modification were summarized in 2.5 kb windows and correlated within each strain (Figure 5A-D). Heterochromatic marks, H3K27me3 and H3K9me3, were highly positively correlated with one another and negatively correlated with H3K27ac. Repressive histone marks were positively correlated with SNPs occurring in TEs, while activating histone mark H3K27ac was associated with SNPs occurring in genes. INDELs followed similar correlations as SNPs (Figure 5A-D). To further understand associations between DNA variation, genomic features, and histone modifications, we tested for independence between histone modifications and variation by genomic feature (see methods for details). The results indicate a significant association between certain DNA variants and histone modifications (Figure 5E, *χ*^2^ critical value of 10.8, Bonferroni multiple testing correction, and Supplemental Tables 2-5). For genes, the *χ*^2^ test indicated that the occurrence of DNA variants was dependent on the presence of all three histone marks across the four genomes (Figure 5E). Variation in TEs mainly occurred independently of histone modifications for both SNPs and INDELs, except for TE variation in O-137 and O-219 associated with H3K9me3 (Figure 5E). For other intergenic regions of the genome, the results indicated that the occurrence of SNPs was dependent on H3K9me3 and to a lesser extent H3K27me3, and both histone marks influenced the occurrence of INDELs (Figure 5E). To understand the direction and nature of the dependence between histone modifications and variation as indicated by the *χ*^2^test, we calculated the frequency of SNPs and INDELs per kb for each genomic feature marked by either H3K27ac, H3K27me3, or H3K9me3 (Table 3 and Table 4). All four genomes showed higher SNPs/kb in genes and intergenic regions when the features were present in heterochromatin domains H3K27me3 or H3K9me3 (Figure 5F). We observed that the SNPs/kb were 1.8 to 4 times higher for genes in H3K27me3 or H3K9me3 domains versus the genes not in the domains (Figure 5F, Table 3). In intergenic regions, SNPs/kb were 1.7 to 3.2 times higher in H3K27me3 domains, and those in H3K9me3 domains were 2.4 to 4.7 times higher compared to regions outside of those domains (Figure 5F, Table 3). Compared to the results for genes and intergenic regions, the SNPs/kb in TEs were contrasting between the heterochromatic histone modifications. The TEs in H3K27me3 domains had on average 50% lower SNPs/kb frequency compared to those not in H3K27me3 domains, while TEs in H3K9me3 domains had on average 3.8 times higher SNPs/kb than TEs not in H3K9me3 domains (Table 3). Given that each comparison within a histone modification used the same set of TEs, the results indicate that TEs in H3K9me3 domains have higher SNP/kb frequency than those in H3K27me3 domains. The ratio of INDELs/kb followed a similar pattern to SNPs/kb. INDELs/kb in genes were 2.4 to 3.3 times higher in H3K27me3 domains and 1.3 to 3.2 times higher in H3K9me3 domains. Intergenic regions have 1.9 to 2.9 times more INDELs/kb in H3K27me3 domains and 1.6 to 3.2 times more in H3K9me3 domains compared to regions outside of those domains. TEs in H3K27me3 domains have on average one third as many INDELs/kb compared to TEs outside of H3K27me3 domains, whereas TEs in H3K9me3 domains have 1.7 to 2 times as many INDELs/kb compared to other TEs. To control for the impact of selection on SNP frequencies, we calculated the frequency of silent SNPs (i.e., SNPs that do not change amino acids) per kb in genes both inside and outside of H3K27me3 domains. On average across all four strains, genes in H3K27me3 domains had a significantly two times higher frequency of silent SNPs/kb compared to genes outside of H3K27me3 domains (Figure 5G, 1.54 versus 0.74 SNPs/kb). These results indicate a clear and statistically significant association between SNP and INDEL frequency and the presence of histone modifications. The exact direction and magnitude differ between some genomic features and histone modifications, but in general both marks associated with heterochromatin were associated with SNPs and INDELs in genes across the four strains. The identification of significantly elevated silent SNPs/kb for genes present in H3K27me3 domains indicates these regions are more mutation-prone.

**Figure 5.**
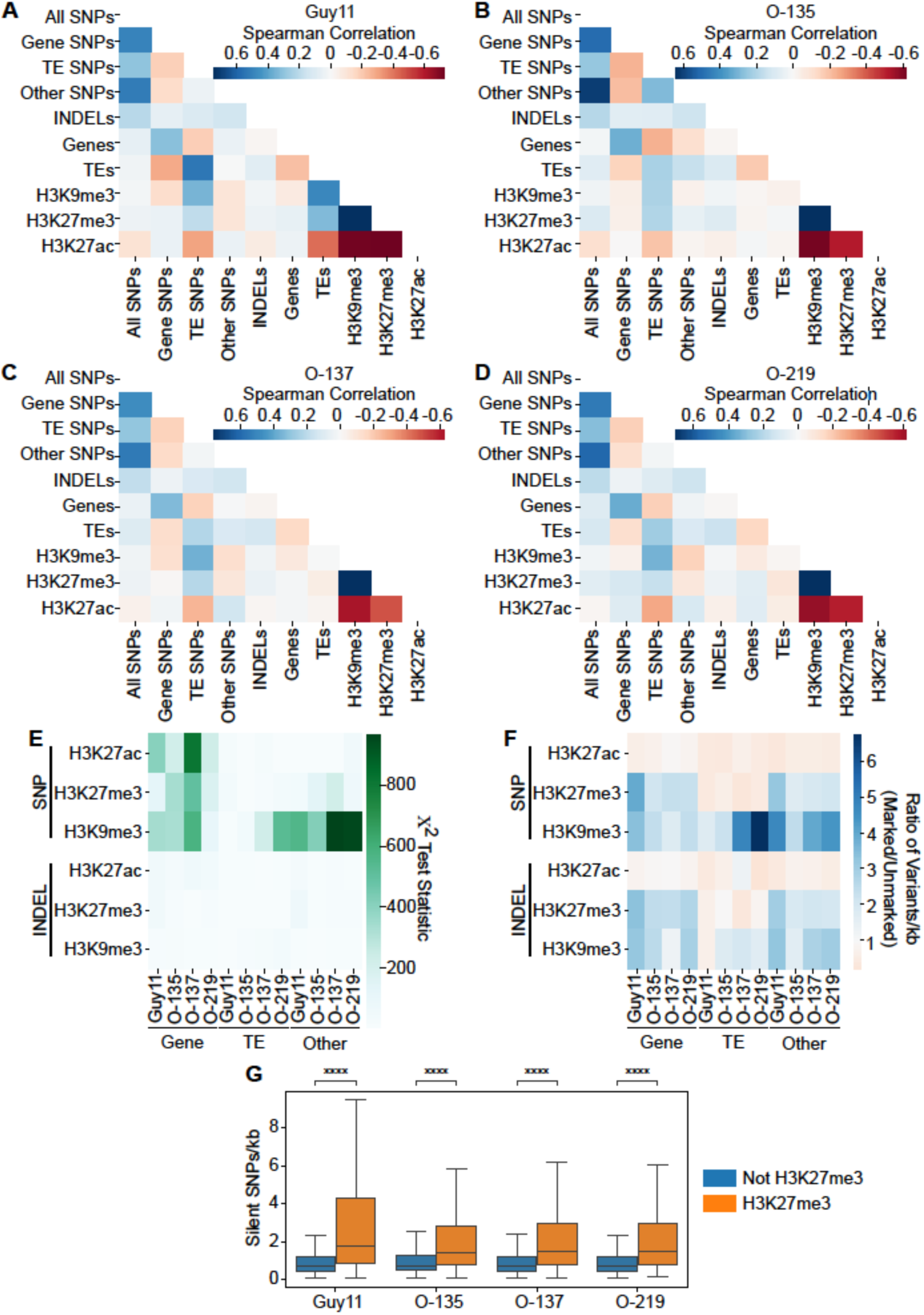
The presence of DNA variation is influenced by histone modifications. (A) Heatmap summarizing Spearman correlation between genes, TEs, SNPs, INDELs, and normalized ChIP-seq read mapping for histone marks H3K9me3, H3K27me3, and H3K27ac in the Guy11 genome. (B) The same as A for strain O-135. (C) The same as A for strain O-137. (D) The same as A for strain O-219. (E) Heatmap summarizing *χ*^2^ results testing independence between the presence of SNPs or INDELs with H3K27ac, H3K27me3, or H3K9me3 in genes, TEs, and intergenic regions. Results from each genome shown on the x-axis. Color bar represents that *χ*^2^ statistic from each test, with a critical *χ*^2^ value of 10.8 at α= 0.001 to reject the null, Bonferroni multiple testing correction. (F) Heatmap depicting the variants per kb as a ratio of marked variants over unmarked variants. Higher values indicate more variants present in marked regions than unmarked regions. (G) Box and whisker plot of silent SNPs/kb across all genes that are either in (orange) or outside (blue) of H3K27me3 domains. Statistical significance detonated as **** for p < 0.0001 as determined by T-test with Bonferroni multiple testing correction.

**Table 3.**
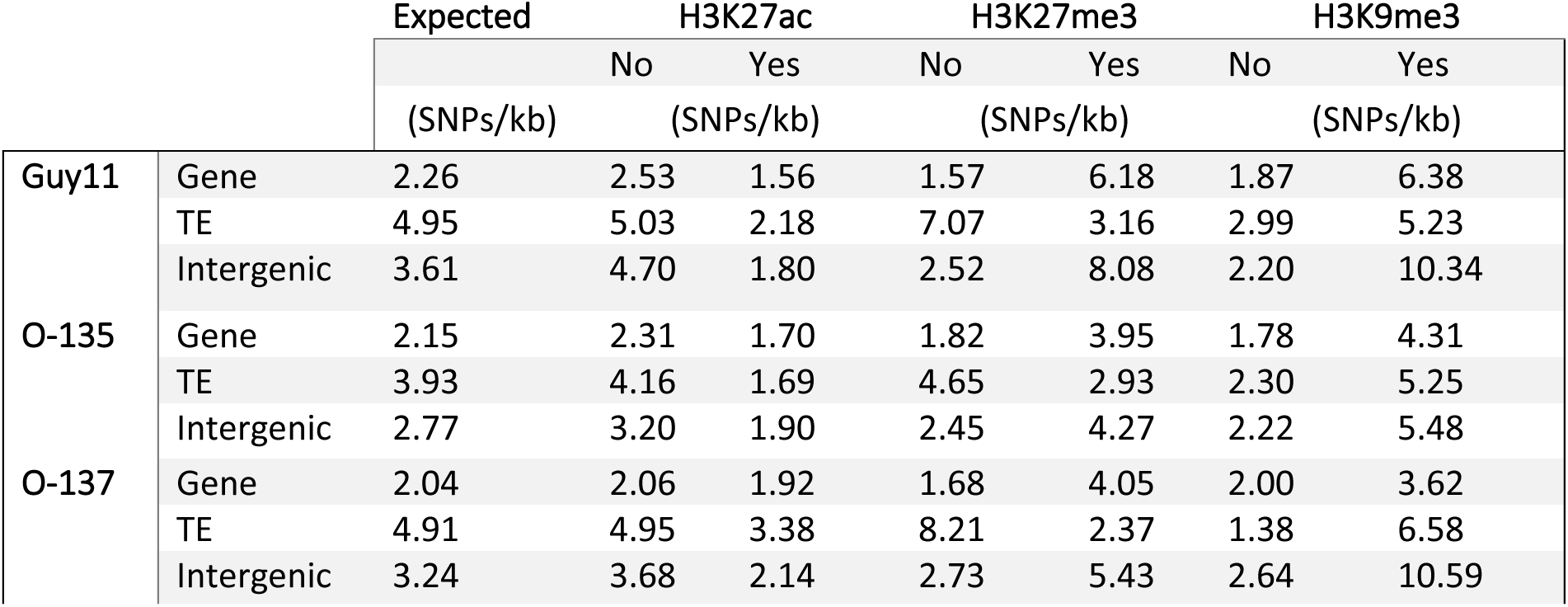
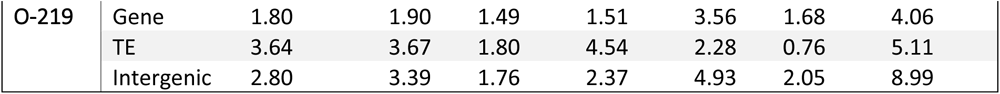
Enrichment of SNPs/kb in genes, TEs, and intergenic regions residing in histone marked regions.

**Table 4.** Enrichment of INDELs/kb in genes, TEs, and intergenic regions residing in histone marked regions.

|  |  | Expected | H3K27ac |  | H3K27me3 |  | H3K9me3 |  |
| --- | --- | --- | --- | --- | --- | --- | --- | --- |
|  |  |  | No | Yes | No | Yes | No | Yes |
|  |  | (INDs/kb) | (INDs/kb) |  | (INDs/kb) |  | (INDs/kb) |  |
| Guy11 | Gene | 0.06 | 0.06 | 0.05 | 0.04 | 0.14 | 0.05 | 0.15 |
|  | TE | 0.06 | 0.06 | 0.03 | 0.08 | 0.05 | 0.09 | 0.06 |
|  | Intergenic | 0.12 | 0.15 | 0.07 | 0.09 | 0.25 | 0.09 | 0.28 |
| O-135 | Gene | 0.05 | 0.05 | 0.05 | 0.04 | 0.10 | 0.04 | 0.11 |
|  | TE | 0.06 | 0.06 | 0.04 | 0.06 | 0.05 | 0.04 | 0.07 |
|  | Intergenic | 0.07 | 0.08 | 0.06 | 0.06 | 0.11 | 0.06 | 0.10 |
| O-137 | Gene | 0.04 | 0.04 | 0.04 | 0.03 | 0.08 | 0.04 | 0.05 |
|  | TE | 0.05 | 0.05 | 0.07 | 0.08 | 0.03 | 0.03 | 0.06 |
|  | Intergenic | 0.09 | 0.10 | 0.07 | 0.08 | 0.16 | 0.08 | 0.23 |
| O-219 | Gene | 0.04 | 0.04 | 0.04 | 0.03 | 0.09 | 0.04 | 0.11 |
|  | TE | 0.04 | 0.04 | 0.01 | 0.04 | 0.04 | 0.03 | 0.05 |
|  | Intergenic | 0.08 | 0.10 | 0.06 | 0.07 | 0.15 | 0.07 | 0.22 |

### Genome synteny is organized by chromatin states

To move beyond pair-wise correlations and gain a better understand of how genome structure influences variation, dimensional reduction through principal component analysis (PCA) was used to integrate all genomic variables studied. For this, genomic data were summarized over 2.5 kb windows, and the genome windows were further divided into syntenic and non-syntenic regions based on whole genome alignments to capture genome variation at a larger scale than the short DNA tracts captured by SNPs and INDELs. Regions were classified as syntenic if at least half the genomic window was shared between the four genomes based on Syri analysis (72), all other windows were classified non-syntenic (Supplemental Table 6). In the Guy11 genome, the first principal component (PC 1) explained 28.6% of the variance, largely separating features associated with euchromatin and heterochromatin (Figure 6A). Gene number, short track DNA variation, and H3K27me3 are the main drivers separating the regions on the second principal component (PC 2), explaining 17.4% of the variance (Figure 6A). It is also clear that genomic windows are separated by synteny, largely along PC 1, even though synteny was not a variable in the PCA. Analyzing short variants by synteny status shows statistically significantly more SNPs and INDELs in non-syntenic genomic windows (Figure 6A). Non-syntenic genomic windows also have significantly fewer genes and more TEs (Figure 6A). Summarizing histone modification ChIP-seq reads by synteny status shows that the euchromatic mark H3K27ac is significantly higher in syntenic regions, while heterochromatic marks H3K9me3 and H3K27me3 were significantly higher in non-syntenic regions (Figure 6B). For the other three genomes, PC 1 separated similar features, namely chromatin features and genome synteny (Figure 6C, E, G). The overall PCA for O-219 looked more similar to Guy11, and O-135 and O-137 looked more similar to each other. The difference was that in O-135 and O-137, genes, effectors and H3K27me3 were less associated with INDELs and SNPs (Figure 6C, E). Summarizing features by synteny, again see that each genome has significantly more SNPs, INDELs, TEs, H3K27me3 and H3K9me3 in non-syntenic regions compared to syntenic (Figure 6D, F, H, and Supplemental Figure 9). These results show that there is conservation between the genomes for organization, with chromatin status helping to define genomic regions that are associated with variation at all levels-SNPs, INDELs, and synteny. The associations are not identical between the genomes, indicating organizational differences, but the effect of chromatin and synteny appears highly similar.

**Figure 6.**
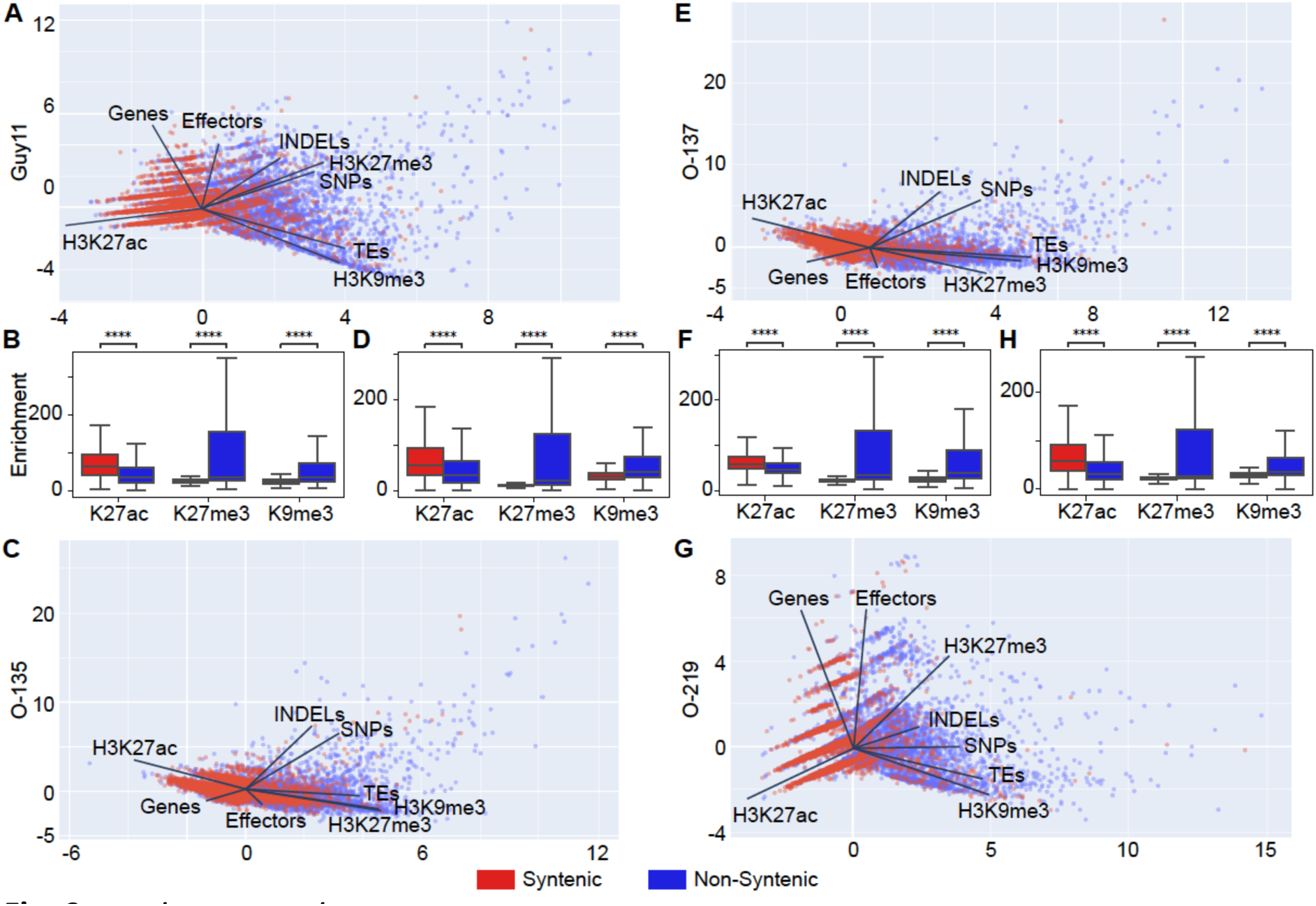
Chromatin clustering in PCA analysis distinguishes syntenic and non-syntenic regions. (A) Principal component analysis (PCA) of summary data for genes, TEs, SNPs, INDELs, H3K27ac, H3K27me3, and H3K9me3 analyzed over 2.5 kb genomic windows of strain Guy11. Red points are windows classified syntenic and blue points are non-syntenic windows. Loadings for each variable are shown as a black vector. (B) Normalized ChIP-seq read mapping in syntenic (red) and non-syntenic (blue) genomic windows for histone modifications H3K27ac, H3K27me3, and H3K9me3 shown as a box and whisker plot for Guy11. (C, D) the same information as show in A, B for strain O-135. (E, F) the same information as show in A, B for strain O-137. (G, H) the same information as show in A, B for strain O-219. Statistical significance detonated as **** for p < 0.0001 as determined by T-test with Bonferroni multiple testing correction.

### Effectors in H3K27me3 domains are more variable in the population and have higher rates of non-synonymous SNPs

A major driver of plant-microbe biology is the function and detection of effector proteins by plants (98). Diversification and deletion of effectors is a major factor in pathogen evolution, influencing virulence, host-range, and niche-adaptation (99, 100). As such, we were interested to determine how the genome wide patterns identified here specifically pertain to effectors. In order to generate the best list of effectors, we searched for conserved sequences in our genomes from a group of 863 predicted effectors recently reported from the strain 70-15 (75). To identify orthologs between strains, sequences of the 863 genes from 70-15 were aligned to each of our four reference assemblies. The majority of 70-15 predicted effectors (96%, 829/863) had alignments in all four strains, of which, 82.1% to 82.5% aligned to a single locus, indicating high single-copy conservation. The remaining 17.9% to 17.5% of predicted effectors aligned to multiple loci, resulting in the total number of aligned loci between 1063 and 1092 (Supp Table 7). Using the identified loci from each of the four genomes, presence/absence variation (PAV) across the 86 rice genomes was calculated based on read mapping coverage. Loci that had read mapping coverage below 65% in 10% or more of the 86 strains were considered to be variable, and the other loci were labeled stable (Table 5). Our previous genome wide analysis indicated that heterochromatic histone marks are associated with DNA variation. To test this further at the gene level, ChIP-seq for H3K27ac, H3K27me3, and H3K9me3 were summarized for each stable and variable predicted effectors (Figure 7A-D). The stable effectors, which showed moderately high read mapping in at least 90% of the strains, had significantly more H3K27ac than variable effectors, while variable effectors had significantly more H3K27me3 and H3K9me3 enrichment (Figure 7A-D). These results are consistent with the genome wide analysis, and further reinforce the idea that H3K27me3 is associated with variation in the genome. In addition to PAV, we analyzed short track DNA variation. For this, we only analyzed effector loci classified as stable to avoid problems with divergent or missing data. As there is a strong selective pressure on many effectors that influences variation in the population, we focused our analysis on silent SNPs that do not result in protein coding changes and are theoretically not impacted by selection. We found that the frequency of silent SNPs/kb was significantly higher for stable effectors located in H3K27me3 marked regions compared to those not covered by H3K27me3 domains in all four strains (Figure 7E-H, 1.45 versus 1.03 SNPs/kb, respectively). These findings indicate that when selection is controlled for, H3K27me3 marked genomic regions accumulate more SNPs in effector coding sequences, which is consistent with the results reported for all genes in the genome.

**Figure 7.**
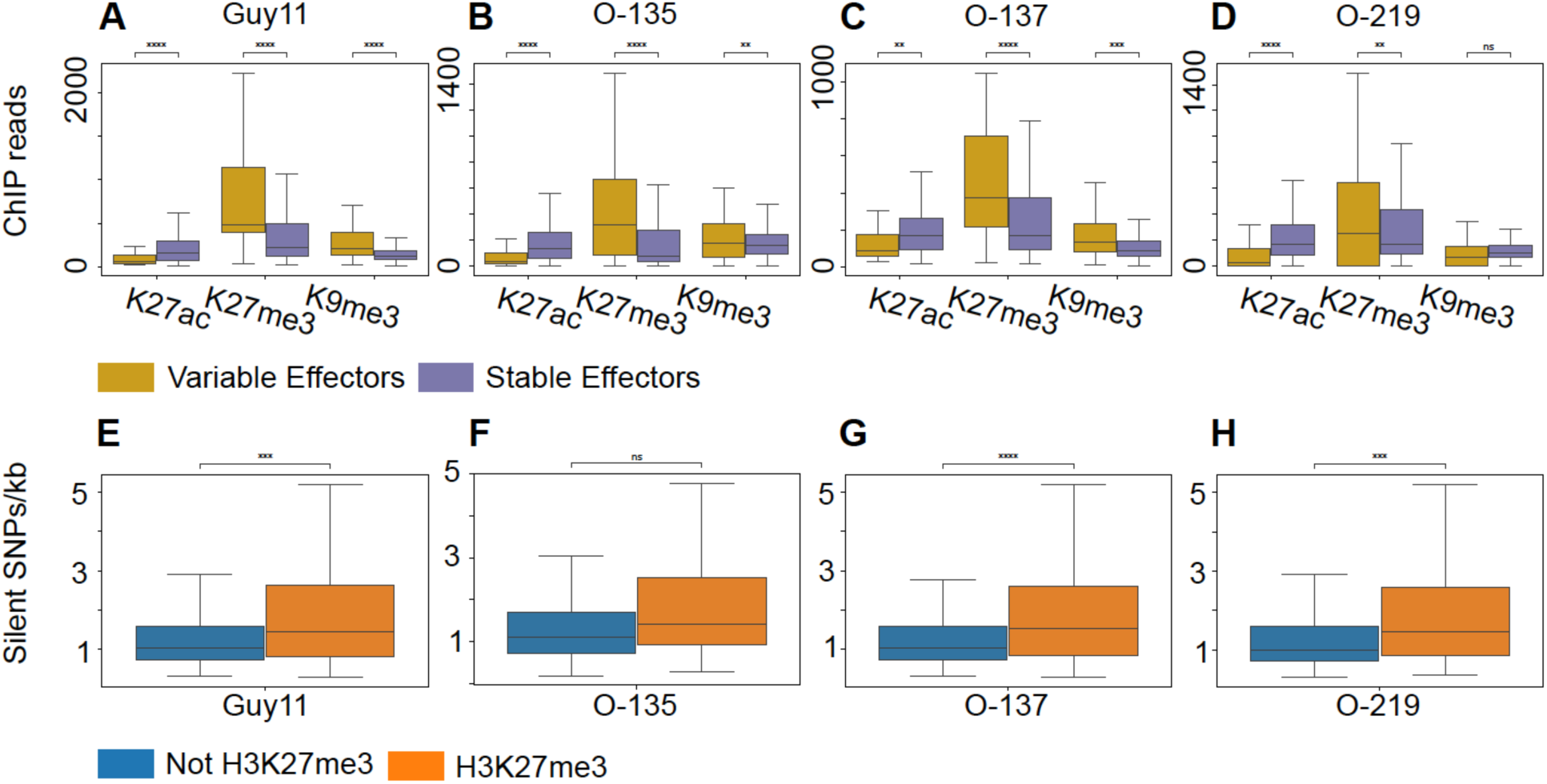
H3K27me3 is associated with variable effectors and a higher frequency of silent SNPs. (A-D) Normalized read mapping of ChIP-seq to stable and variable of effectors of modifications H3K27ac, H3K27me3, and H3K9me3. (A) Guy11, (B) O-135, (C) O-137, (D) O-219 (E-H) Silent SNPs/kb frequency of stable effectors that reside inside or outside of H3K27me3 marked regions. (E) Guy11, (F) O-135, (G) O-137, (H) O-219. Statistical significance detonated as ns = not significant, ** p < 0.01, *** p < 0.001, **** p < 0.0001 as determined by T-test with Bonferroni multiple testing correction.

**Table 5.** Stable and variable predicted effector genes.

|  | Stable | Variable |
| --- | --- | --- |
| Guy11 | 995 | 70 |
| O-135 | 981 | 92 |
| O-137 | 1013 | 50 |
| O-219 | 1002 | 77 |

## DISCUSSION

This research tested the hypothesis that DNA variation is associated with physical and epigenetic characteristics of the genome. Previous inquiries into the connections between DNA variation and genome organization have described the two-speed genome composed of rapidly evolving and more stable genomic regions of plant pathogenic microbes (8, 101, 102). In the *M. oryzae* genome, there is no evidence that gene-density is associated with variability or evolutionary signal as original described in *P. infestans* (8). Instead, the *M. oryzae* genome is organized by epigenetic histone marks onto a continuum of stable euchromatin regions and more variable heterochromatic regions. These results are based on the analysis of short track DNA variation, namely SNPs and INDELS, as well as genome level synteny and gene level PAV, from high-quality genome assemblies and histone modification maps. This research further describes the characteristics of genome regions with contrasting DNA variability, and gene density does not seem to be a defining characteristic. Integrated analysis of histone modifications indicates that the epigenome is a more significant factor in defining stable and hyper-variable regions of the genome.

Analysis of the four *M. oryzae* genomes shows clear evidence of similar organization. The four strains examined in this study represent three divergent lineages of rice-infecting *M. oryzae*, but they have largely similar associations between histone modifications and DNA variation. Effector genes, TEs, SNPs, and INDELs are unevenly distributed throughout the genome, with most localizing to sub-telomeric regions of chromosomes. Likewise, the location of SNPs and INDELs correlates with the occurrence of H3K27me3 and H3K9me3. This organization is consistent with the description of genome compartments that may facilitate effector diversification (99). SNPs in particular show a bimodal distribution of dense and sparse SNPs, and dense SNPs occur much more frequently in regions occupied by repressive histone marks, H3K27me3 and H3K9me3. An argument could be made that the high occurrence of dense SNPs is a byproduct of heterochromatin and associated TEs (94, 103). However, on average across the genomes, 67% of dense SNPs are present in genes or intergenic regions, indicating that TEs do not drive the dense SNP association with repressive histone marks. Also, the frequency of SNPs in H3K27me3 marked genes is higher than in unmarked genes, again showing the association is not TE dependent. For H3K9me3 marked features, SNPs are more frequent regardless of whether those features were genes, TEs, or intergenic regions. Similar to SNPs, INDELs are more frequently found in genes and intergenic regions marked with H3K27me3 and H3K9me3, while TEs that contained INDELs occurred in H3K9me3 domains. It was also remarkable to find that H3K27me3 is associated with non-syntenic regions, effectors that are variably present in the population, and even non-synonymous SNPs. One notable difference between the four strains was evident in the PCA representations that showed a different magnitude of association between SNPs, INDELs and H3K27me3. The results indicate that the O-135 and O-137 genomes are overall more similar to one another despite the strains coming from two different asexual lineages of *M. oryzae* (41, 82). This may be related to their mode of reproduction, but the exact cause and consequence remains unknown.

It is currently unclear whether epigenetic marks lead to specific types of variation in the genome or whether it is a response to DNA variation. Experimental evolution in *Z. tritici* revealed that TEs had significantly higher base substitution mutation rates when the sole methyltransferase responsible for H3K9me3 (*Kmt1*) was deleted (32, 33). This is a clear demonstration of the canonical role of heterochromatin and H3K9me3 providing genome stability and defense against TEs (104). However, in a mutation accumulation study in *N. crassa*, the mutation rate for DNA in H3K9me3 marked regions was 10-fold higher compared to euchromatic DNA (36). Likewise, in both *Z. tritici* and *N. crassa*, there was a clear, albeit moderate, increase in mutation accumulation in regions marked by H3K27me3 (33, 36). It seems that these histone modifications, and likely others, play a balancing role in repressing DNA templated functions, while either imperfectly maintaining DNA fidelity or actively promoting change. Direct measures of DNA fragility, such as the occurrence of DNA breaks, are not well documented in filamentous fungi, but in *N. crassa*, the localization of phosphorylated H2A, termed γH2A, a hallmark of DNA damage, localizes to heterochromatin (105, 106). Therefore, the creation of DNA damage and DNA repair may be an important component in creating variation. In eukaryotes, the types of mutations as well as kinetics of repair enzymes have been reported to change between euchromatin and heterochromatin (35, 107). Therefore, a potential mechanisms for the observed differences between the associations of SNPs and INDELs with histone marks could be that the different chromatin states influence DNA repair pathway choice in a given region, and thus alter the frequency and type of mutations (108). However, these associations are confounded by the tendency for TEs and effectors to be in subtelomeric regions of chromosomes, which are documented to be fragile due to highly repetitive sequences (109). Further studies to isolate and test specific factors controlling the frequency and spectrum of mutations are needed to understand cause and effect (7). As such, it remains unclear whether the presence of effectors helps drive variability or if effectors arise in regions that are predisposed to variability due to genome organization. Our data on the frequency of silent SNPs/bp shows that when controlling for selection, genes in H3K27me3 marked regions have accumulated more silent mutations than genes outside of H3K27me3 regions. This was true for all genes in the four genomes, and also when looking at the specific case of effectors. This suggests that H3K27me3 marked regions have experienced higher rates of mutation in the four *M. oryzae* genomes given that the silent SNPs should be generally free from selection. Further understanding the interplay between the epigenome and the creation of variation will have a profound impact on our understanding of evolution.

## METHODS

### Genome assemblies, repeat and gene annotation

High-molecular-weight DNA was extracted following the protocol from (43). DNA > 30 kb were further enriched by BluePippin size selection. Oxford Nanopore sequencing was performed using MinION sequencing platform. Raw MinION fast5 files were processed into fastq files by Guppy (version 3.4.4) (44) with the following parameters: --disable_pings --compress_fastq --flowcell FLO-MIN106 --kit SQK-LSK109. Adaptors were trimmed from reads by Porechop (version 0.2.4) (45). Canu (version 1.9 for O-135 and 2.2.1 for O-137, O-219 and Guy11) (46) was used for genome assemblies with the following parameters: genomeSize=45m, corMhapSensitivity=normal, corOutCoverage=80, minOverlapLength=2000, minReadLength=5000, correctedErrorRate=0.12, useGrid=1. The draft assemblies were further polished by Nanopolish (two times, version 0.13.2) (47) and Pilon (two times, version 1.22) (48), with Nanopore and Illumina reads. Each contigs were renamed based on the NUCmer (version 3.1) (49) alignment with the core chromosomes from B71ref2 and 70-15 genome assemblies (50, 51). Unplaced contigs with (>50%) duplicated with the core chromosomes or bacterial DNA were manually removed from the assemblies. The size of mitochondrial DNA was trimmed according to B71ref2 assembly.

Predicted results for repeated sequences in the four isolates from LTRfinder (version 1.07) (52), LTRharvest (version 1.5.10) (53), MGEscan (version 3.0.0) (54), LTR_retriever (version 2.9.0) (55), MITE-Tracker (version 2.7.1) (56) and MITE-Hunter (57) were combined with CD-HIT (version 4.8.1) (58), parameters: -c 0.8 -G 0.8 -s 0.85 -T 15 -aL 0.85 -aS 0.85, to remove highly similar sequences. The non-redundant library for repeated sequences were used as input for annotation with Repeatmasker (version 4.0.9) (59).

The soft-masked genome was used as the input for gene annotation following funannotate pipeline (version1.8.5) (60). The completeness of assembled genomes and predicted gene annotations were evaluated by BUSCO (version 5.1.2) (61) by using ascomycota_odb10 dataset. The presumed secreted proteins were predicted by SignalP (version 5.0ba) (62) and further annotated with EffectorP (version 3) (63). This research project only used DNA sequences from the seven core chromosomes for analysis. The mini chromosomes and unplaced contigs are included in the assembly, and were annotated, but these regions were excluded from this analysis to focus on the core chromosomes found in all strains of *M. oryzae*.

### CHEF-gel preparation and electrophoresis

To prepare the CHEF-gel plug, 1×10^8^ concentrated fungal protoplasts (37 μL) (64) were mixed with 2% CleanCut Agarose (63 μL, Bio-Rad, #1703594). The mixture was then loaded into a plug mold (Bio-Rad, #1703713). The solidified plugs were treated in NDS buffer (65) containing 400 μg/mL proteinase K (Thermo scientific, #EO0491) at 50 °C overnight, followed by 4x washes with 50mM EDTA. The washed plugs were loaded in 1% Certified Megabase Agarose (2.4g agarose/240mL 0.5x TBE) (Bio-Rad, #1613108). CHEF-gel electrophoresis was performed on a CHEF Mapper System (Bio-Rad) using the following parameters: “Int. Sw. Tm=120s, Fin. Sw. Tm=3,600s, Gradient=2.0 v/cm, Run time=96h, Included Angle=120°, Temperature=14°C”. The size of the mini-chromosome was measured using *H. wingei* Ladder (Bio-Rad, #1703667).

### Chromatin Immunoprecipitation Sequencing and analysis

Chromatin Immunoprecipitation sequencing was performed following our previously published methodology (22). Briefly, proteins were crosslinked to DNA in mycelia grown in CM liquid culture using formaldehyde and then flash frozen in liquid nitrogen. 200 mg of frozen tissue per sample was ground and DNA was sheared using sonication. Sheared DNA was treated with MNase and then precleared using an 80:20 ratio of A and G beads. Samples were then incubated for 14 hours with the following antibodies: H3K27ac (Diagenode, C15410196), H3K27me3 (abcam, 6002), H3K36me3 (abcam, 9050), H3K4me3 (Diagenode, C15410003), or H3K9me3 (Diagenode C15410056). Antibodies were then pulled down with the A/G beads and washed thoroughly. DNA was purified using the iPure kit from Diagenode. Libraries were prepared using NEBNext FFPE DNA repair mix and Ultra II End-prep mix. A universal adaptor was then ligated to DNA samples and IDT i5 and i7 primers were used in a PCR reaction to add unique barcodes to each of the samples. DNA fragments were double size-selected after PCR to enrich for approximately 300 bp size libraries. Sequencing was performed using Illumina NextSeq high output, single end sequencing. Peak calling was performed using MACS2 (66).

### Single nucleotide polymorphism detection

Single nucleotide polymorphisms (SNPs) were detected in the 86 rice-infecting strains using the Genome Analysis Toolkit (GATK v3.4.4) (67). Each of the 86 strains was mapped against each of the four strains (Guy11, O-135, O-137, and O-219) using gatk MarkDuplicates, then indexed and HaplotypeCaller was run using -ploidy 1 -ERC GVCF. VCF files were combined using gatk CombineGVCFs and genotyped using gatk GenotypeGVCFs. Then variants with low data to support them were removed using gatk VariantFiltration –filterexpression “AN < 68” and vcftools –gzvcf –remove-filtered-all –recode (68) to remove filtered variants. AN < 68 was chosen so that only variants that had sequencing data, either for the reference or the variant, in at least 80% of the 86 strains were used in further analysis. VCF entries with more than one variant at a position were removed so that all variants were biallelic using gatk SelectVariants – restrict-alleles-to BIALLELIC. SNPs were extracted from the list of variants using gatk SelectVariants –select-type-to-include SNP. Finally, each remaining SNP was kept only if at least one of the strains in which a SNP was called only had reads mapping to the alternative allele and none supporting the reference. SNPs only supported by mixed read mapping strains were removed from the analysis.

SnpEff (69) was used to identify the impact of SNPs and determine whether they were synonymous or nonsynonymous. Custom databases were made using GFF files from our genome assemblies. Biallelic high-confidence filtered VCF files were used as input files to generate the SNP calls.

### INDEL detection

Insertion and deletion (INDEL) variations were detected in the 86 rice-infecting strains using the Genome Analysis Toolkit (GATK v3.4.4) (67). Each of the 86 strains was mapped against each of the four strains (Guy11, O-135, O-137, and O-219) using gatk MarkDuplicates, then indexed and HaplotypeCaller was run using -ploidy 1 -ERC GVCF. VCF files were combined using gatk CombineGVCFs and genotyped using gatk GenotypeGVCFs. Then variants with low data to support them were removed using gatk VariantFiltration –filterexpression “AN < 68” and vcftools –gzvcf –remove-filtered-all –recode (68) to remove filtered variants. AN < 68 was chosen so that only variants that had sequencing data, either for the reference or the variant, in at least 80% of the 86 strains were used in further analysis. INDELs were extracted from the list of variants using gatk SelectVariants –select-type-to-include INDEL. VCF entries with more than one INDEL found at a reference position were removed so that all variants were biallelic using gatk SelectVariants –restrict-alleles-to BIALLELIC. Finally, each remaining INDEL was kept only if at least one of the strains in which an INDEL was called only had reads mapping to the alternative allele and none supporting the reference. INDELs only supported by mixed read mapping strains were removed from the analysis.

### Average pairwise distance between the strains

Filtered VCF files of *M. oryzae* isolates aligned to assemblies were split by chromosome. Single chromosome variant files were converted into LDHat (2.2) (70) input files. Once the sites and loci input files were generated by chromosome, using a custom perl script to split the chromosome into sliding windows of 5 kb total, stepping 2.5 kb each step, we calculated genetic diversity statistics by window. Average pairwise distance was normalized by window size to generate the nucleotide diversity stat on a per site basis. Window estimates for diversity statistics were plotted along the chromosome and outliers were generated based on while genome distribution.

### Synteny Analysis

To analyze synteny between the four genomes, a pipeline was used to map the genome assemblies to each other and then extract syntenic regions. Assembled genomes were mapped in a one-to-one fashion using Minimap2 (71). With the following options ‘-x asm5’ preset option for comparison between related genomes with less than 5% divergence. The output alignment files were parsed using the Syri program with default parameters (72). To identify syntenic regions between the four genomes, syntenic regions were intersected using bedtools intersect to keep only regions that are syntenic between all four strains (73). Finally, the syntenic information was integrated into the 2.5kb windows for each genome. For this, a 2.5kb window was classified as syntenic if at least half of the window overlapped a syntenic region. All 2.5kb windows not classified as syntenic from this analysis were classified as non-syntenic.

### Integrated Genomic Analysis

The expected frequency of SNPs or INDELs per kb was calculated by dividing the total number of variants, SNPs or INDELs, over the total size of a gene, TE, or intergenic region. Then the number of variants in features was split whether it was in a marked (H3K27ac, H3K27me3, H3K9me3) region, previously determined using MACS2, and divided by the size of those sub-groups. Deviation between expected and observed variants was determined using a chi-square analysis, chi2_contingency (74). Bonferroni multiple testing correction was performed to adjust the critical value to 10.8 at df = 1. Principal component analysis was performed on the data subdivided into 2.5 kb windows. Genomes features were counted within these windows and ChIP enrichment was mapped into 2.5 kb windows. K-means clustering was performed on the first two components. The distribution of SNPs and INDELs in these clusters was measured.

### Predicted Effector Gene Analysis

Predicted effector genes from strain 70-15 were gathered from available data from (75). Gene IDs were used to obtain protein sequences from Ensembl fungi (76). Protein sequences were aligned to our four strains through tBLASTn (77) using custom nucleotide databases constructed from our genome assemblies. Alignments were filtered to remove alignments with an e-value greater than 0.1 and with a percent ID less than 50%, then alignments within 2 kb were merged to connect multiple exons of a gene into one alignment, then alignments that were less than half of the size of the protein sequence were removed to filter out small common peptide sequences. Read coverage at each of the alignment loci was calculated using Illumina reads of the 86 rice-infecting strains. Loci where 10% or more of the strains had less than 65% read mapping coverage of the region were considered to be variable loci while all other loci were considered to be stable.

## Data availability

Genomic sequencing and assemblies for strains Guy11, O-137, 0-135, and 0-219 have been deposited in the National Center for Biotechnology Information (NCBI) Sequence Read Archive (SRA) under Bioproject accession PRJNA1006359. The BioSamples are SAMN37051403 (Guy11), SAMN37099169 (O-135), SAMN37099170 (O-137), and SAMN37099171 (O-219). Data for Guy11 ChIP-seq using anti-H3K27me3, anti-H3K27ac, and associated input were collected previously and accessed through NCBI SRA Bioproject accession PRJNA646251.

## Code availability

The python code used for analysis, along with intermediate summary files, and gene and transposable element annotations, have been deposited in the Zenodo digital repository https://doi.org/10.5281/zenodo.8233842.

## Funding

This work was supported by the National Science Foundation (NSF) through the Models for Uncovering Rules and Unexpected Phenomena in Biological Systems (MODULUS) (award no. MCB-1936800) to DEC, United State Department of Agriculture-National Institute of Food and Agriculture (USDA-NIFA) (award no. 2018-67013-28492) to DEC, USDA NIFA award (2021-67013-35724) to SL, BV, and DEC, and the NSF award (2011500) to SL, BV, and DEC. The funders had no role in study design, data collection and analysis, decision to publish, or preparation of the manuscript. Mention of tradenames or commercial products in this publication is solely for the purpose of providing specific information and does not imply recommendation or endorsement by the U. S. Department of Agriculture. USDA is an equal opportunity provider and employer.

## Conflict of Interest Disclosure

The authors declare no competing interests.

---

**Supplemental Figure 1.**
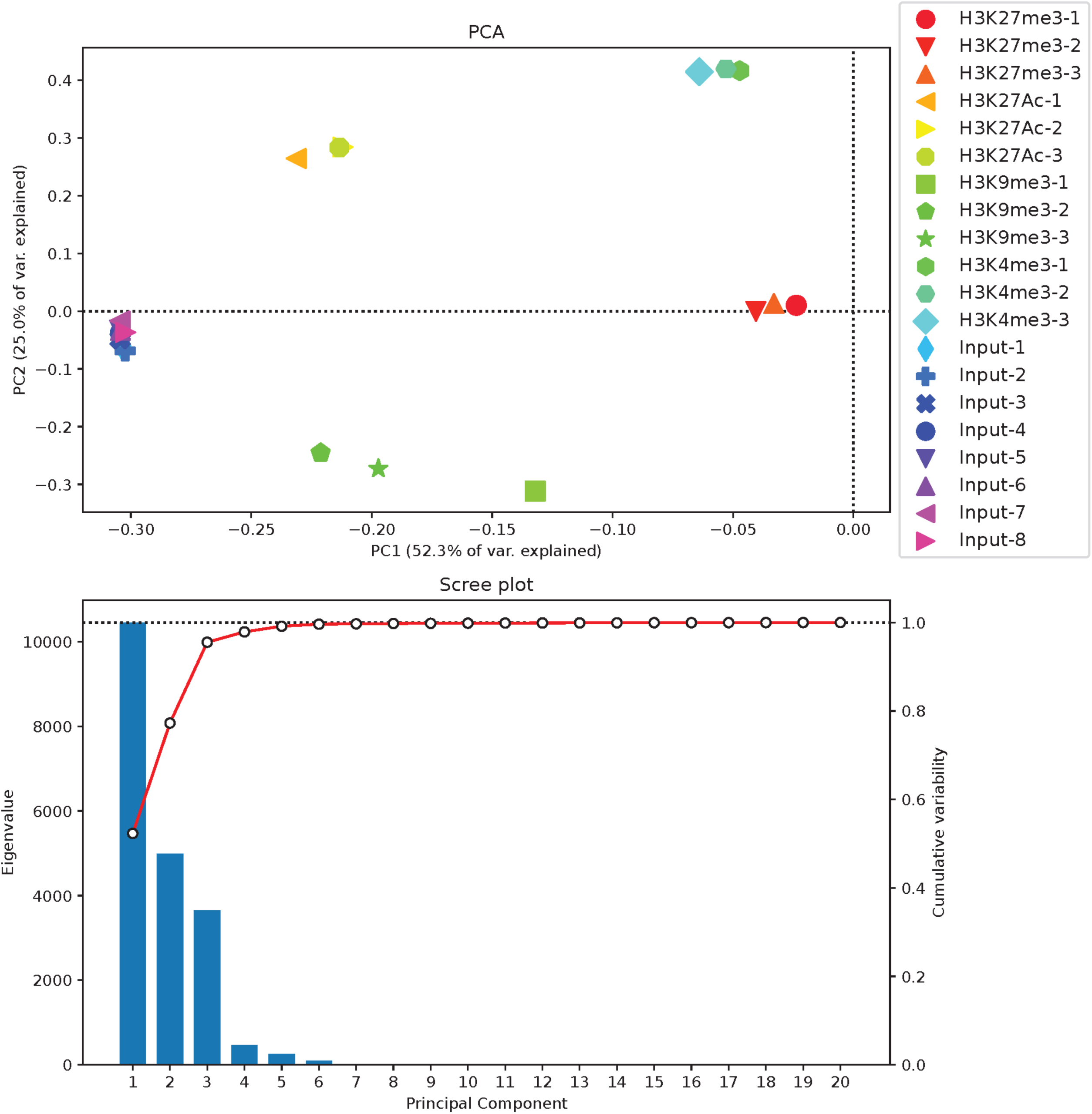
Guy11 Chromatin Immunoprecipitation PCA. Principal component analysis of individual replicates of chromatin immunoprecipitation sequencing in Guy11 for the epigenetic histone marks H3K27me3, H3K27ac, H3K9me3, and H3K4me3. The highest contributing components explain 52.3% and 25.0% of variance.

**Supplemental Figure 2.**
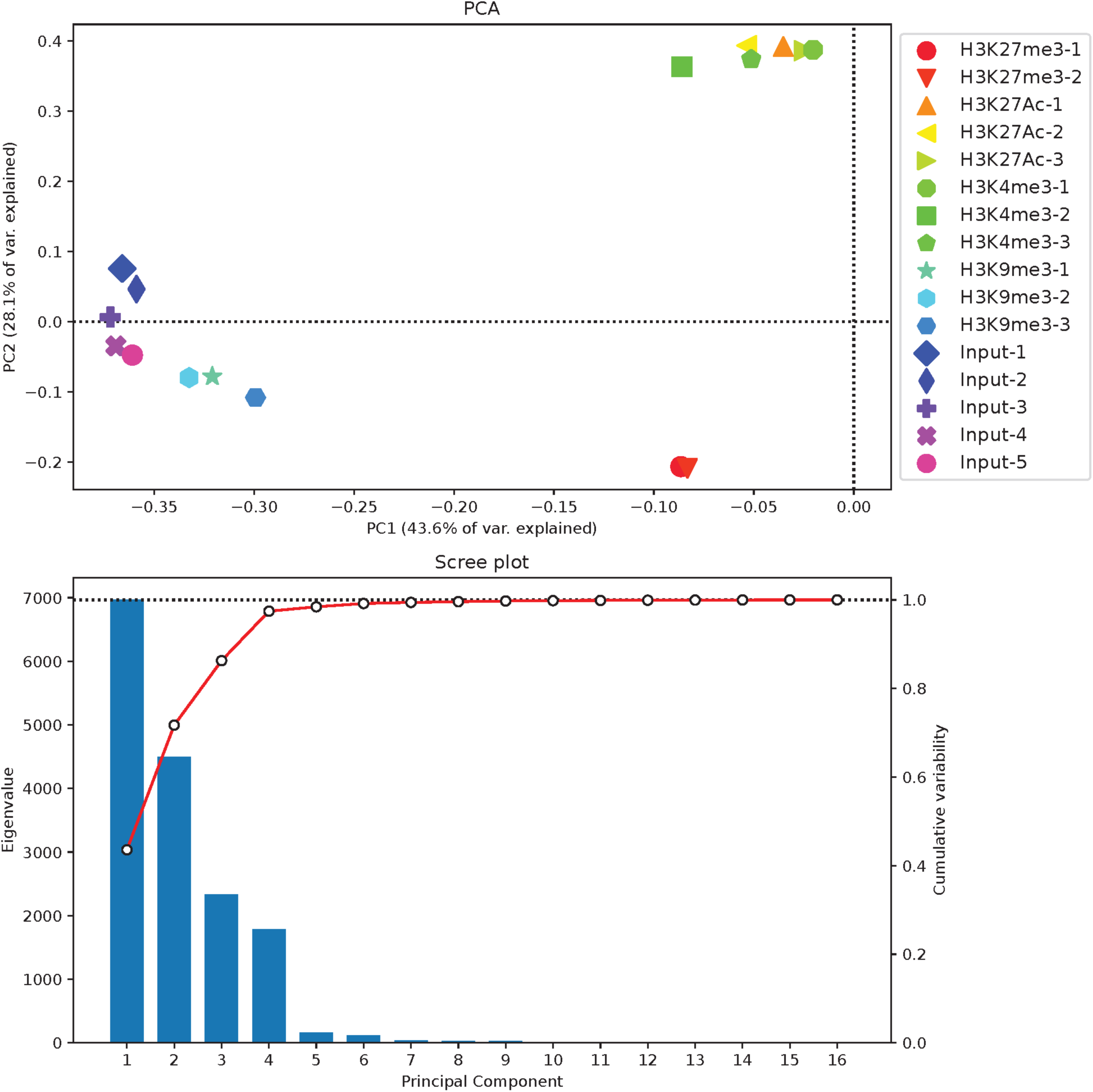
O-135 Chromatin Immunoprecipitation PCA. Principal component analysis of individual replicates of chromatin immunoprecipitation sequencing in O-135 for the epigenetic histone marks H3K27me3, H3K27ac, H3K9me3, and H3K4me3. The highest contributing components explain 43.6% and 28.1% of variance.

**Supplemental Figure 3.**
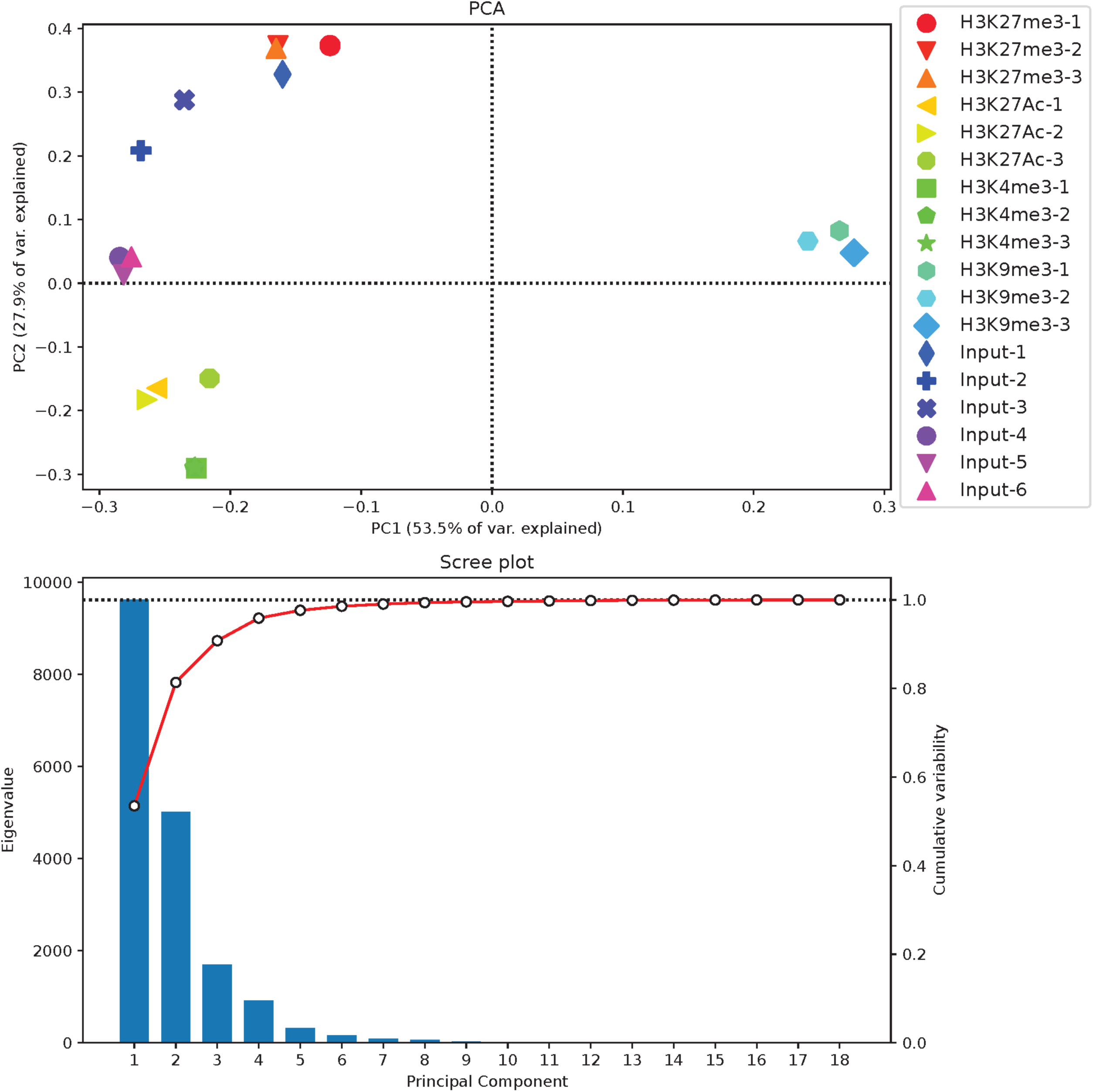
O-137 Chromatin Immunoprecipitation PCA. Principal component analysis of individual replicates of chromatin immunoprecipitation sequencing in O-137 for the epigenetic histone marks H3K27me3, H3K27ac, H3K9me3, and H3K4me3. The highest contributing components explain 53.5% and 27.9% of variance.

**Supplemental Figure 4.**
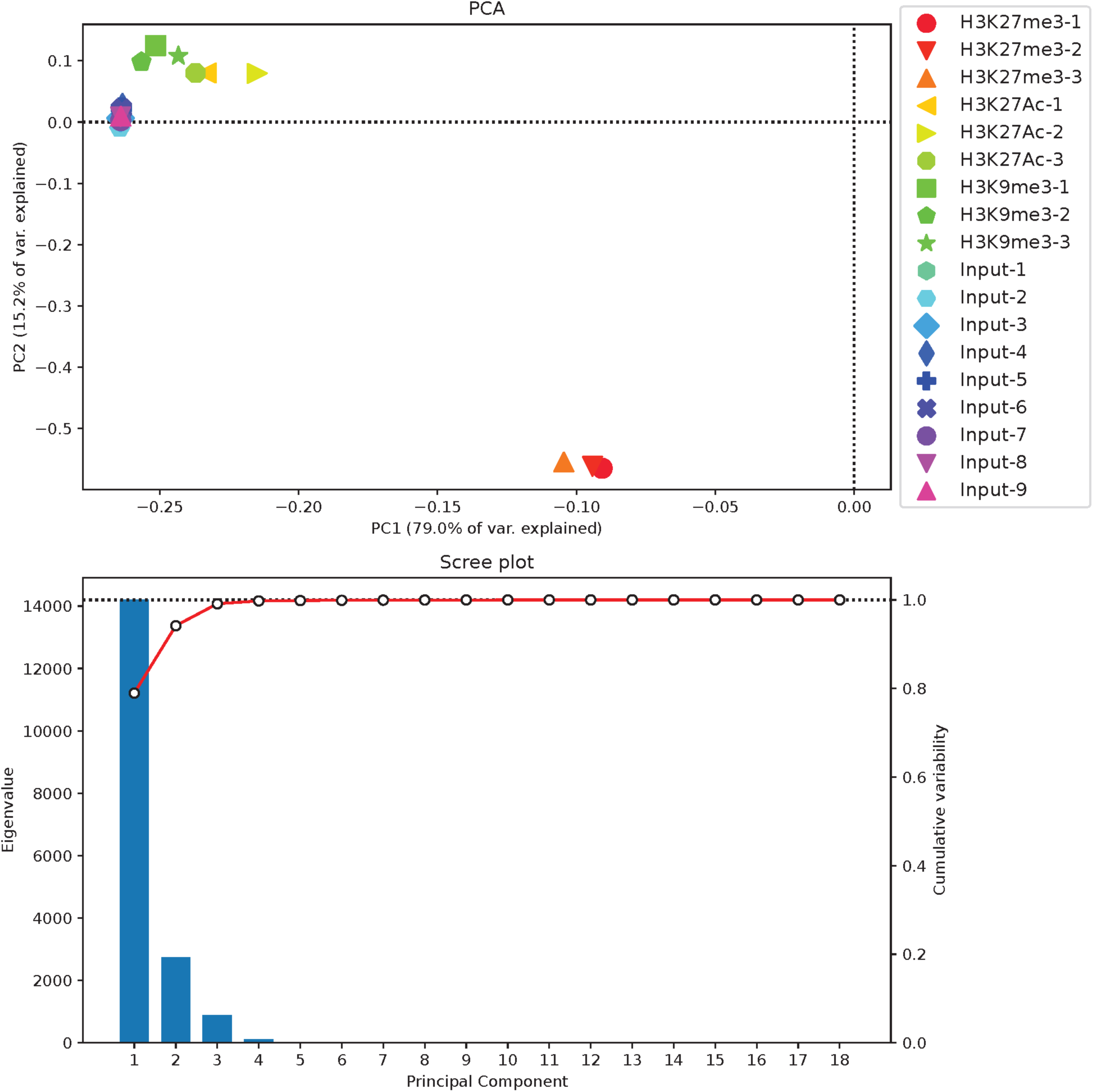
O-219 Chromatin Immunoprecipitation PCA. Principal component analysis of individual replicates of chromatin immunoprecipitation sequencing in O-219 for the epigenetic histone marks H3K27me3, H3K27ac, and H3K9me3. The highest contributing components explain 79.0% and 15.2% of variance.

**Supplemental Figure 5.**
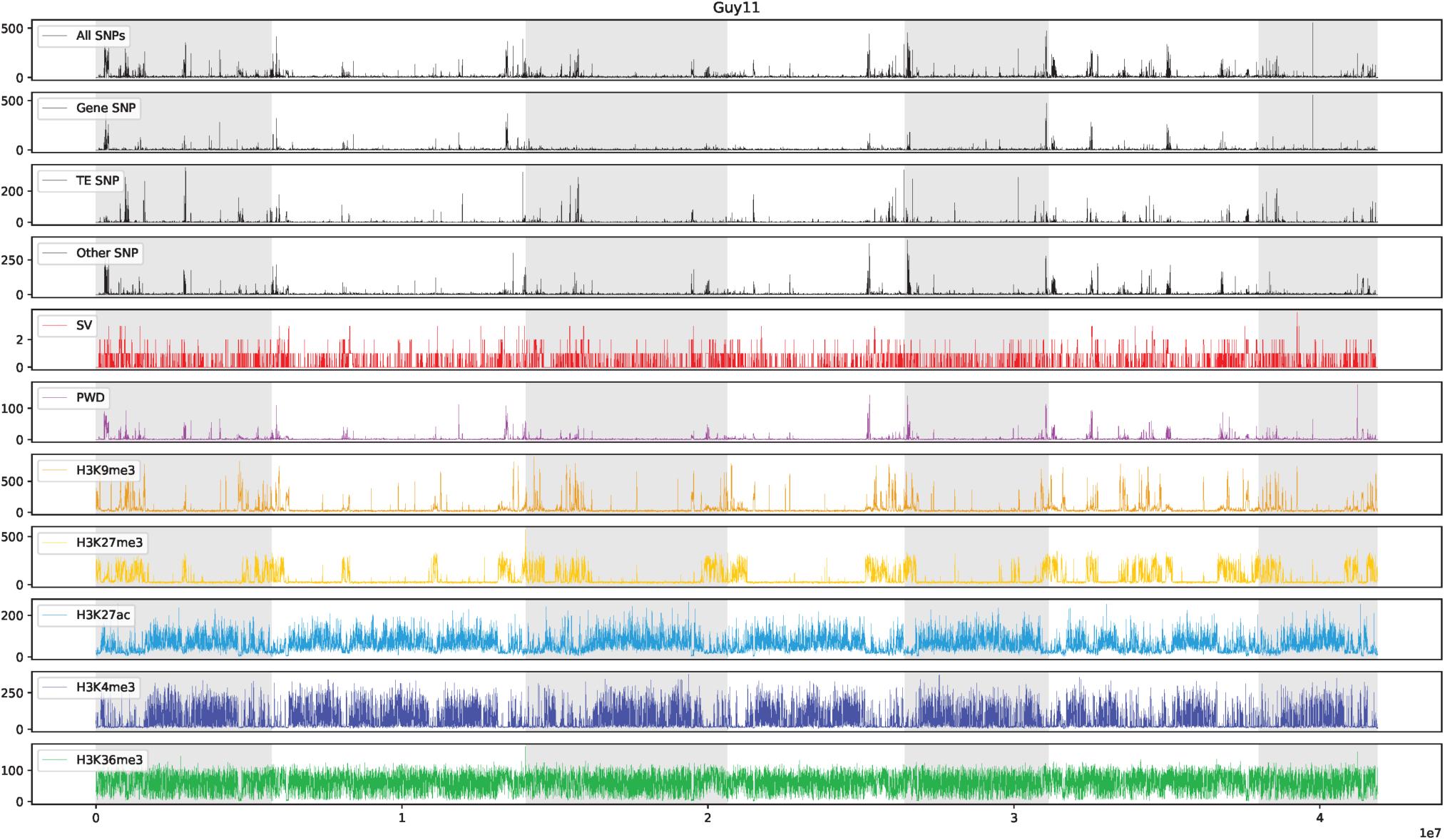
Guy11 whole genome mapping of variation and histone marks. Graph of SNPs, INDELs, average pair-wise distance (PWD) and ChIP enrichment of H3K9me3, H3K27me3, H3K27ac, H3K4me3, and H3K36me3. Alternating shaded regions indicate core chromosomes 1 through 7.

**Supplemental Figure 6.**
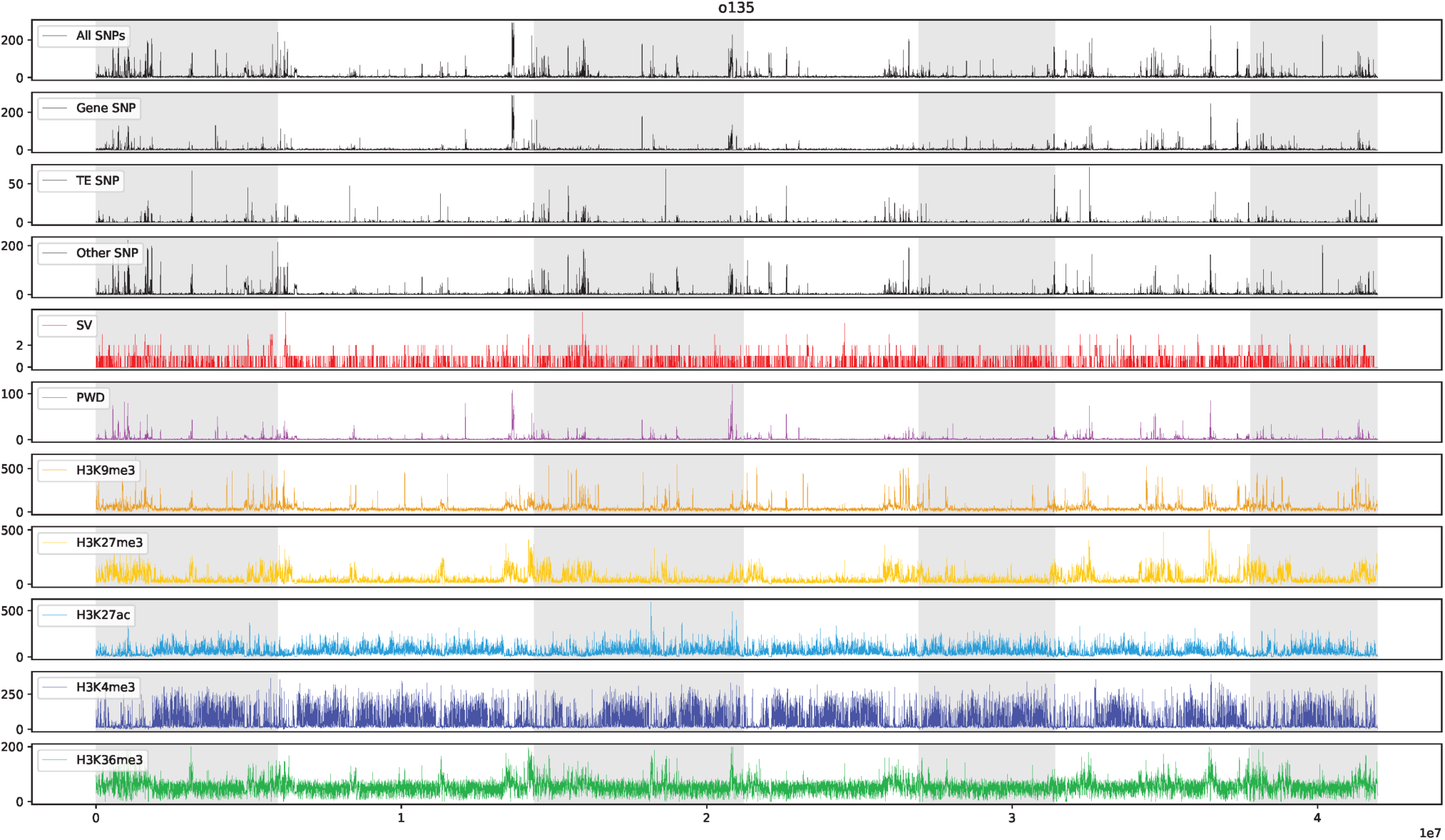
O-135 whole genome mapping of variation and histone marks. Graph of SNPs, INDELs, average pair-wise distance (PWD) and ChIP enrichment of H3K9me3, H3K27me3, H3K27ac, H3K4me3, and H3K36me3. Alternating shaded regions indicate core chromosomes 1 through 7.

**Supplemental Figure 7.**
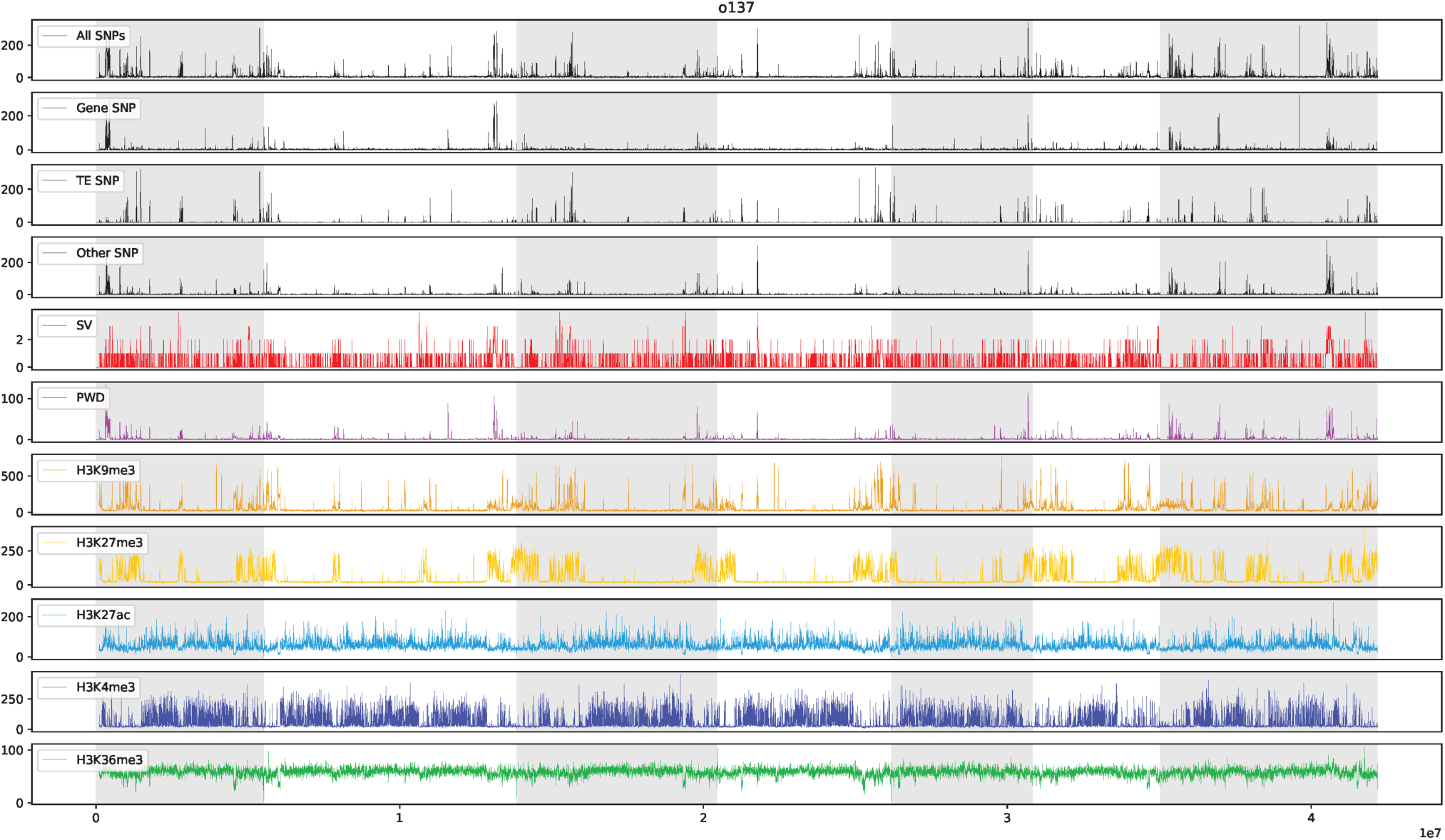
O-137 whole genome mapping of variation and histone marks. Graph of SNPs, INDELs, average pair-wise distance (PWD) and ChIP enrichment of H3K9me3, H3K27me3, H3K27ac, H3K4me3, and H3K36me3. Alternating shaded regions indicate core chromosomes 1 through 7.

**Supplemental Figure 8.**
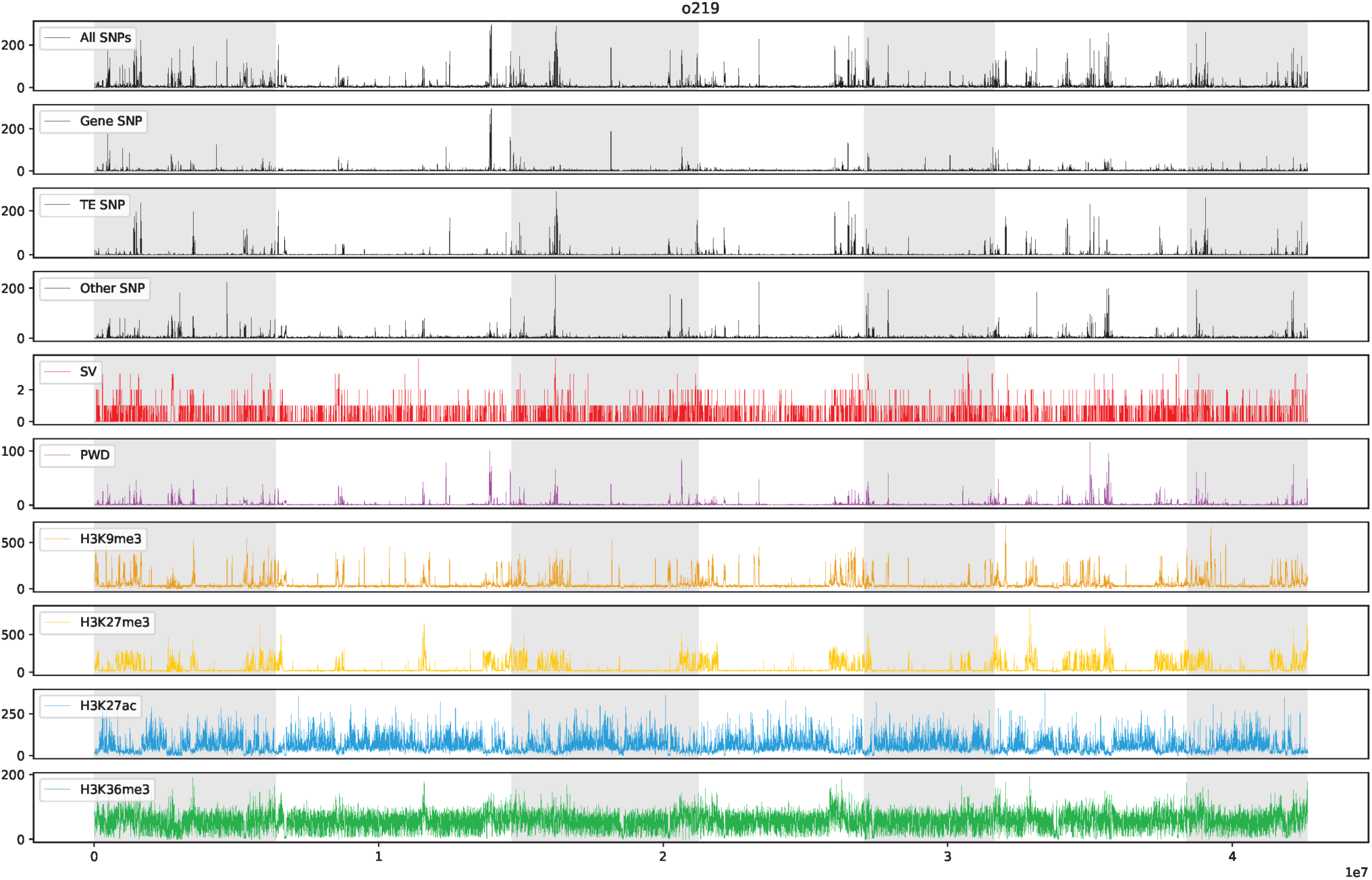
O-219 whole genome mapping of variation and histone marks. Graph of SNPs, INDELs, average pair-wise distance (PWD) and ChIP enrichment of H3K9me3, H3K27me3, H3K27ac, and H3K36me3. Alternating shaded regions indicate core chromosomes 1 through 7.

**Supplemental Figure 9.**
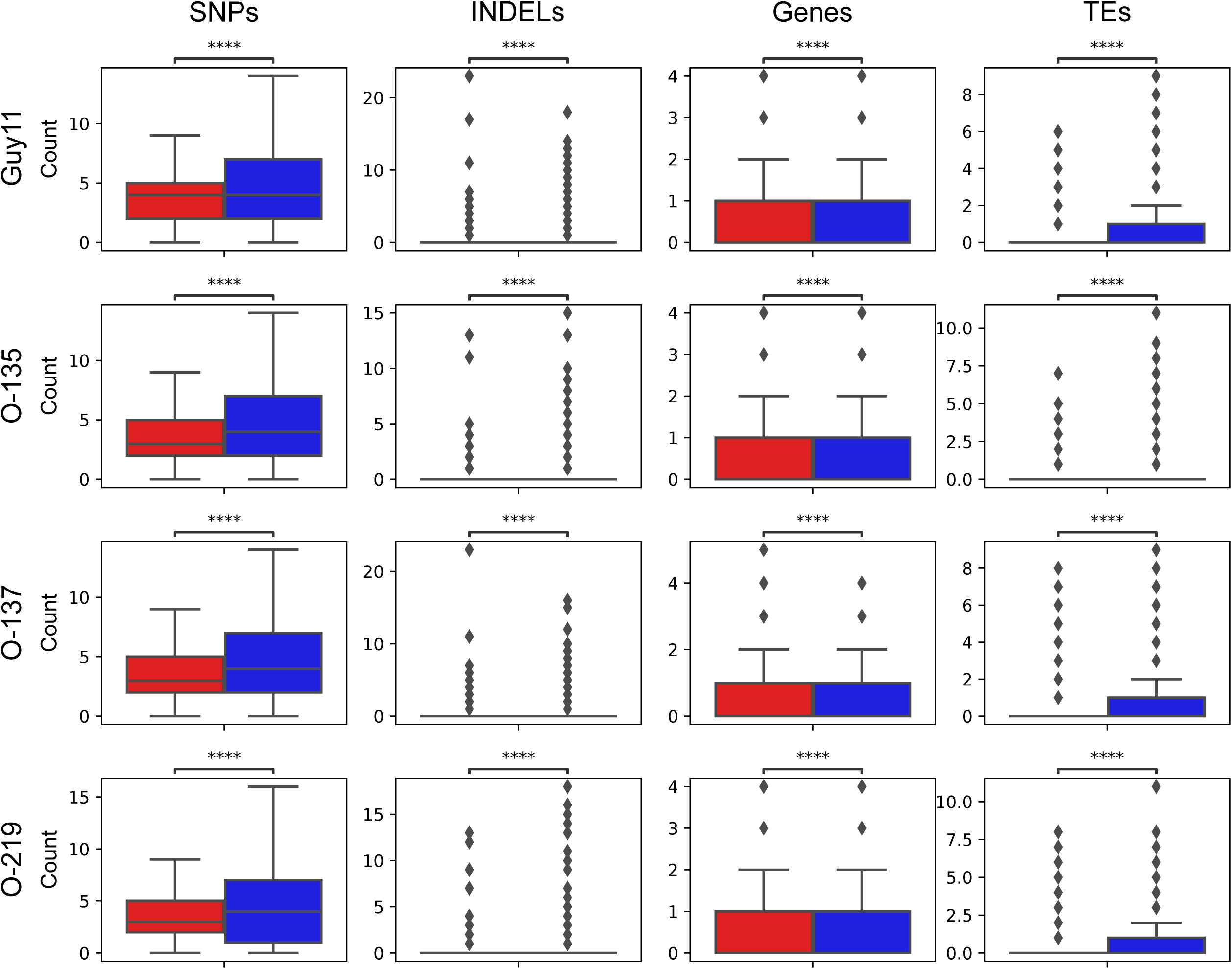
Genomic features associated with syntenic and non-syntenic regions. Box ad whisker plot summarizing the number of SNPs, INDELs, Genes, and TEs present in the 2.5 kb genomic windows, split between syntenic (red) and non-syntenic (blue) regions. Each of the four strains is labeled to the left of each row.

## Notes

### Competing Interest Statement

The authors have declared no competing interest.

